# Alpha-Band Inter-brain Connectivity during Minimal Tactile Interaction: An Exploratory Hyperscanning Study

**DOI:** 10.1101/2025.06.18.660494

**Authors:** Chen Lam Loh, Leonardo Zapata-Fonseca, Mark M. James, Guillaume Dumas, Tom Froese

## Abstract

Researchers have begun to examine the minimal conditions required for inter-brain coupling (IBC) in humans, and brain-to-brain synchrony can emerge during technology-mediated interactions, not only in shared physical spaces. Here, we present an exploratory electroencephalography (EEG) hyperscanning study investigating whether inter-brain synchronization (IBS) can emerge under conditions of both spatial isolation and minimal interaction. We used the Perceptual Crossing Experiment (PCE), a real-time task in which paired individuals search for one another in a one-dimensional virtual tactile environment using only minimal haptic cues. Dyad-shuffled surrogate analysis identified a small set of candidate inter-brain connections, with the strongest observed connections in the alpha bands. Unsupervised clustering identified four recurrent behavioral patterns and three inter-brain connectivity patterns. Descriptively, the two less common inter-brain configurations occurred more frequently during phases characterized by greater inter-avatar separation and reduced contact, although this association was not supported after accounting for the nested structure of the data. While preliminary, these findings provide a proof-of-concept framework for relating neural synchronization to interactive behavior in a highly constrained setting. The candidate connections were not individually significant, and whether inter-brain coupling arises under such conditions remains to be tested in hypothesis-driven designs.

## Introduction

Hyperscanning, the simultaneous neuroimaging of more than one individual, has become a standard method to study intra- and inter-brain neurodynamics during real-time social interaction [12]. Because terminology varies, we use inter-brain synchronization (IBS) for the phase synchronization of neural signals between individuals [16, 47], and inter-brain coupling (IBC) as a general term for their association, including amplitude correlation [60]. IBC has been proposed to facilitate information transmission between brains, improving the prediction of others’ actions and supporting more complex forms of joint behavior [27]. Using electroencephalography (EEG) hyperscanning, IBS is found to occur in different frequency bands and brain regions, most commonly in the alpha and beta frequency bands [16, 23, 35, 43, 61, 64, 67, 69, 70], and involve the prefrontal cortex (PFC) and right temporo-parietal junction (rTPJ) [12, 19, 37, 66].

Most hyperscanning experiments to date have relied on visual or auditory interaction channels, for instance, in speech and communication tasks [34, 36, 61], musical performance tasks [54, 63], and cooperative joint action tasks [4, 67, 69]. In contrast, touch is a more fundamental means of establishing social presence [51], and humans possess a sophisticated capacity for tactile sensorimotor control that enables exploration, object manipulation, and social bonding through physical contact [53]. Notably, social touch alone has been associated with alpha-band IBC in the somatosensory (frontal and central) regions and is important for the communication of empathy and social understanding [23]. However, prior touch-based hyperscanning studies typically involved face-to-face contact or shared physical environments [23, 56], leaving open the question of whether robust IBC can occur with touch alone when people are physically separated. To address this, here we explore IBC during a purely tactile, minimal social interaction paradigm: the Perceptual Crossing Experiment (PCE) [5, 6, 14, 21, 29, 41, 71, 72].

We define minimal interaction as an exchange that is deliberately constrained in both sensory modality and communicative bandwidth. In the PCE, participants received only haptic feedback within a one-dimensional virtual space, with no access to visual or auditory cues. In our implementation, partners sat physically isolated with eyes closed, each controlling a simple avatar in the shared 1D virtual environment via a hand-held interface. Their task was to cooperatively locate each other within this virtual space. Although the PCE includes simple avatars and objects, the overall design deliberately minimizes the information available for coordination. Crucially, participants were not given details about the task structure before the experiment. To succeed, they had to actively develop novel interaction strategies in real time. This reflects the idea that communication and coordination can emerge from the dynamics of interaction itself, even under highly constrained conditions [27].

What remains unclear is how such severely constrained conditions affect inter-brain dynamics. Recent evidence suggests that simply sharing an environment may be a key requirement for IBC: for instance, significant IBC has been observed when two people interact in proximity [42, 50], and even when interacting in a virtual (though not physically isolated) environment [25]. Conversely, IBC tends to diminish in purely online video interactions with no shared environment [64], although there are reports of IBC between physically separated participants under certain tasks [69]. Thus, the PCE offers a test case for whether engaging in a minimally shared virtual tactile environment can elicit inter-brain coupling. If we observe IBC under these conditions, it would suggest that even minimal touch-based interaction is sufficient to induce IBS despite physical isolation.

Using a publicly available PCE dataset released by our group [44], we take an exploratory, descriptive approach with two aims: to examine whether IBS emerges in any frequency band during minimal touch-based interaction and to explore how behavioral variables relate to inter-brain measures. Rather than testing specific mechanistic hypotheses, we present a framework for jointly analyzing behavioral dynamics and neural coupling in restricted virtual environments, illustrating how hyperscanning can be applied to minimal social interactions and generating observations for future hypothesis-driven work.

## Materials and Methods

### Sample Description

Sixty-four participants (thirty-two dyads) took part in the ECSU-PCE study [44]. The sample included individuals from 24 nationalities who could understand and communicate in English. Participants’ ages ranged from 18 to 68 years old (mean: 28.6 ± 8.9). Of the 64 participants, 39 identified as female. Most participants were blinded before the experiment and did not know with whom they would interact. In three dyads this was not possible: dyad 32 were classmates, and dyads 15 and 25 were couples. Out of the 32 dyads, dyad 31 had to be disregarded from the analysis due to technical difficulties during the EEG recordings, and dyad 14 was excluded because automated artefact rejection during pre-processing removed 51.7% of its epochs, above the 50% threshold applied to the combined resting and task epochs. The analyses therefore use 30 dyads, that is 60 participants. The experiment was approved by the OIST Human Subject Research Review Committee (HSR-2023-013), and all participants gave informed consent.

### Task Procedure and Data Acquisition

The task was based on the PCE, a two-person interaction model used in embodied cognitive science to study aspects of social cognition such as shared attention, mutual recognition, and the detection of social situations [5, 6, 14, 21, 29, 41, 71, 72]. In the PCE, dyads were seated in two separate but similar rooms (Fig 1). Each player interacted through a haptic interface, the Perceptual Crossing Device, with two hand-held components (one for rotation, one with a button), each containing an integrated vibration motor [18]. With this interface, each player controlled the movements of an avatar along the perimeter of a circle, which served as a shared virtual space. In this space each player could encounter three objects: the other player’s avatar, a static object, and a shadow that mimicked the other avatar at a fixed offset. Overlapping with an object activated that player’s vibration motor in an all-or-nothing manner.

**Fig 1.**
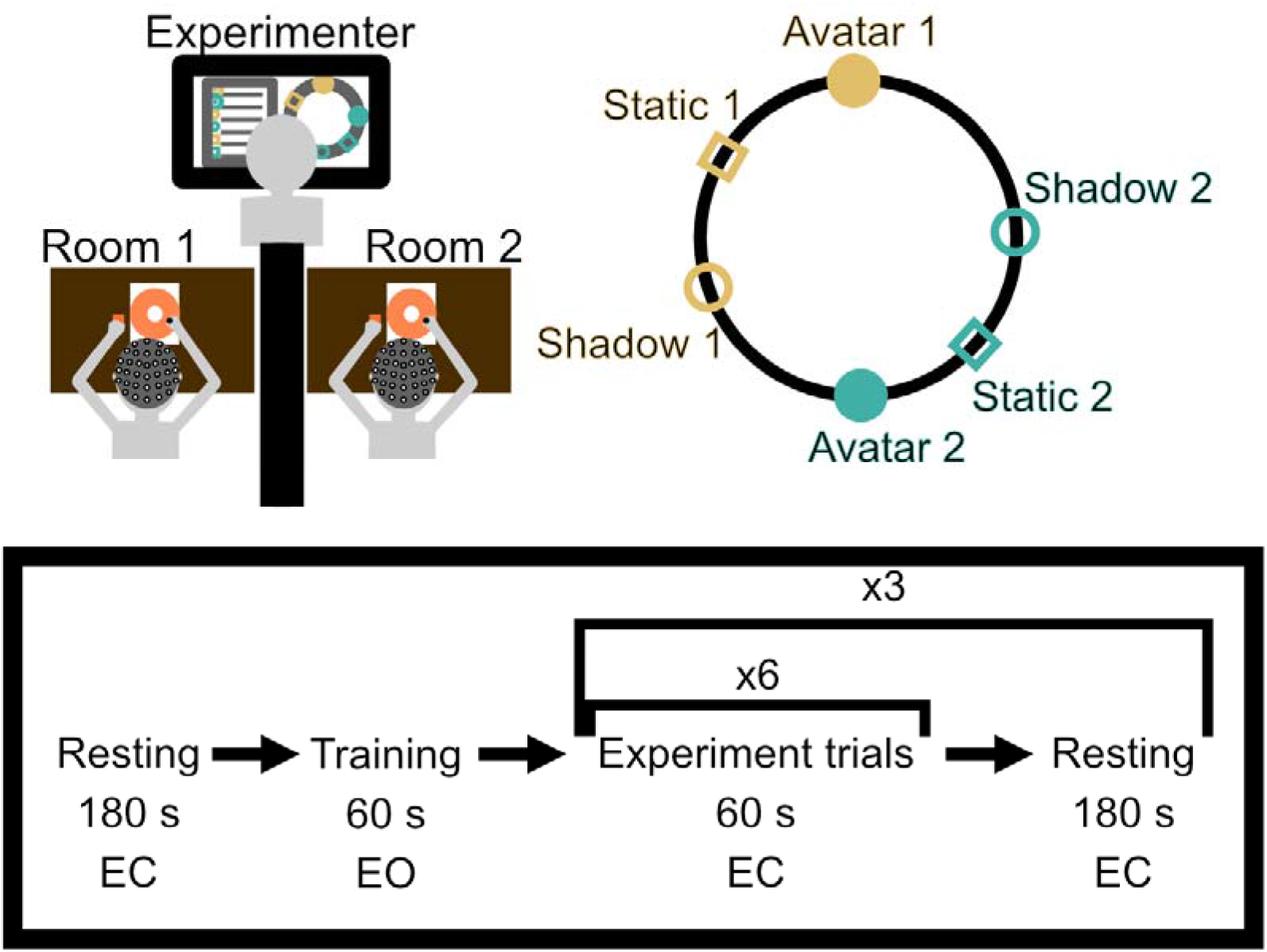
The Perceptual Crossing Experiment (PCE). The top of the figure shows two participants interacting in two different rooms in the minimally-shared space using the haptic human-computer interface. The bottom of the figure shows the experimental timeline. EC=eyes closed; EO=eyes open.

Players interacted within the circular space to identify each other through the vibration patterns, pressing a button whenever they felt the crossed object was the other player’s avatar. Each player was permitted only one button press per trial, although they were encouraged to continue interacting after making their response. The experiment consisted of four eye-closed resting sessions (REST) and eighteen eye-closed task trials (TRIAL). Each REST lasted 180 s, whereas each TRIAL lasted 60 s. Aside from the haptic vibration triggered by object overlap, no feedback regarding task performance or accuracy was provided during or after the trials. All participants completed the trials in the same order specified by the standard PCE protocol.

Following the original study, dual-EEG was simultaneously and continuously recorded with 64-channel BrainAmp amplifiers (Brain Products, Germany). Participants were fitted with a standard 64-channel Brain Product actiCAP snap set with sensors arranged in the international 10-10 system. The low- and high-cutoff frequencies were 0.016 Hz and 1000 Hz, respectively. The ground electrode was Fpz, and the common reference was FCz. The impedance was maintained below 25 kΩ. The sampling frequency was 1000 Hz, and an online notch filter was applied at 60 Hz. Data were recorded with BrainVision Recorder 1.20.0502 (Brain Products).

### Behavioral Data Analysis

The analysis pipeline is summarized in S1 Fig. Using the behavioral dataset, we derived seven behavioral variables. The first three were the distance between the participants’ avatars (P1avatar-P2avatar), the mean avatar–shadow distance (mean of P1avatar-P2shadow and P2avatar-P1shadow), and the mean avatar–static-object distance (mean of P1avatar-P1static and P2avatar-P2static). The next two were the absolute differences in avatar velocity and acceleration between participants (P1vel-P2vel and P1acc-P2acc). These differences were corrected for circular wrapping and averaged within each 3-s epoch during the task. The remaining two variables were the proportion of mutual contact duration (p_mutual_contact) and the dynamic time warping (DTW) [62] distance between participants’ avatar velocities. For p_mutual_contact, the proportion of time during which both vibration motors were active was computed for each 3-s epoch. The DTW computation is detailed in Supplementary Methods S1. Finally, all seven behavioral variables were normalized to the range [0, 1]. An example of the behavioral variables is shown in S2 Fig.

The behavioral variables for each 3 s epoch, dyad, and trial formed a (DYADxTRIALxEPOCHxN) matrix, where N was the number of variables. We applied k-means clustering [48] to these data using the MATLAB kmeans function with 20 replicates to identify behavioral patterns during the PCE. The optimal number of clusters was then determined automatically using the elbow method (see Supplementary Methods S1). The proportions of occurrence were calculated by summing the occurrences of each identified cluster over epochs, trials, or dyads.

The joint success (JS) rate was defined as the total number of joint recognitions [44] of each other’s avatar across all trials of each dyad, and normalized by the total number of trials per dyad.

### EEG Data Analysis

#### Pre-processing of EEG data

We pre-processed the EEG using the Hyperscanning Python Pipeline (HyPyP) [7] with MNE-Python [24], keeping the original 1000 Hz sampling rate. We labeled the data, split it into the two subjects, and applied a zero-phase finite impulse response (FIR) filter from 1 to 100 Hz (5 Hz transition bandwidth at the high cutoff, low cutoff set to auto). After average re-referencing, we segmented the data into 3 s epochs, yielding 20 epochs per task trial and 60 per resting phase. We used 3 s epochs because shorter epochs can give spurious and unreliable estimates of inter-brain coupling [73]. We then applied Infomax independent component analysis (ICA) with 15 principal components. Components classified by ICLabel [45] as non-brain sources (eye, muscle, heart, or noise) were automatically removed using an ICLabel-based procedure adapted from the HyPyP tutorial, with no manual component rejection. We cleaned the epochs with prep.AR_local. Across the 30 dyads, 348.0 of 360 task epochs were retained (*SD* = 12.1, median = 351, range 307 to 360) and 215.7 of 240 resting epochs (*SD* = 33.9, median = 228, range 78 to 240), a mean removal of 3.3% of task epochs (*SD* = 3.3%) and 10.1% of resting epochs (*SD* = 14.1%), leaving 10,440 task and 6,470 resting epochs. Details are in the hypyp_preprocess.py script and S1 Fig. Rest–trial spectral context is shown in S5 Fig (relative Welch power spectral density) and S6 Fig (cluster-permutation t-statistics).

#### Measure of Inter-brain Synchronization

We quantified IBS as phase synchronization between two EEG sensors, computed as the adjusted circular correlation coefficient (CCorr) [73]. The original CCorr measures the covariance between two phase signals and is robust against spurious synchrony [11]. Because EEG signals lack a well-defined mean phase across epochs, the adjusted CCorr treats the phases as uniformly distributed, which stabilizes the estimate [73]. We used the absolute value of CCorr, constraining it to [0,1], because our analyses focused on the magnitude rather than the direction of synchronization. The full definition, following Zimmermann and colleagues [73], is given in Supplementary Methods S2.1. For the rest of the article, CCorr refers to the adjusted CCorr. CCorr was computed via HyPyP for all sensor pairs in all bands of interest. To obtain the analytic signal, zero-phase FIR band-pass filters with Hamming windows and automatic order and transition bandwidth were applied before the Hilbert transform. The frequency bands were Theta (4-7.5 Hz), Alpha-Low (7.5-10 Hz), Alpha-High (10-12.5 Hz), Beta-Low (12.5-20 Hz), Beta-High (20-30 Hz), Gamma-Low (30-58 Hz), and Gamma-High (62-100 Hz), based on prior work linking these frequency ranges to IBS and social interaction [9, 16, 35, 69, 70]. We subdivided the Alpha, Beta, and Gamma ranges into Low and High bands to provide finer frequency resolution while accommodating potential heterogeneity in band-specific activity across dyads [2, 31]. The Alpha subdivision follows EEG research suggesting distinct contributions of lower alpha (attentional and sensorimotor engagement) and higher alpha (top-down inhibition and cognitive control) [39, 40]. Given the exploratory aim, we use these subdivisions for finer resolution but interpret sub-band differences cautiously, without assuming strict dissociations.

#### Inter-brain Functional Connectivity

Because participants shared similar task environments and stimuli in the PCE, we constructed a dyad-shuffled null distribution to distinguish spurious inter-brain correlations from those arising from genuine interaction. Observed circular correlations (CCorr) were evaluated against this null distribution, and only connections that exceeded the surrogate threshold were retained for further analysis. Full details of the null distribution construction, thresholding, and clustering procedures are provided in the Supplementary Methods S2.2.

#### Regions of Interest

For the averaging of sensor-sensor FCs into ROI-ROI FCs, we grouped all 64 sensors into 9 pre-defined ROIs containing five to eight sensors each. The ROIs were defined as right frontal (RF, 7 sensors), left frontal (LF, 7), right central (RC, 5), left central (LC, 5), right temporal (RT, 8), left temporal (LT, 8), right parietal (RP, 8), left parietal (LP, 8), and a midline chain (MID, 8). Each sensor belongs to exactly one ROI. Grouping sensors into predefined ROIs improved interpretability, reduced the number of comparisons, and could improve signal-to-noise ratio. Following previous research [43, 64], we separated the ROIs into left and right ROIs to investigate whether there was a lateralization in inter-brain FC [15, 64]. The sensors included in each ROI are shown in S4 Fig.

### Statistical Methods

The Spearman correlation coefficient between the behavioral variables and the Alpha-Low and Alpha-High ROI-ROI connections was calculated using the corr function in MATLAB. The function provided a *p*-value based on testing against a null hypothesis of no correlation between the variables. For large samples, this is done by approximating the ranked-sum statistic as a t-statistic distribution.

Each epoch was assigned a behavioral cluster and an inter-brain cluster. We then counted the occurrences of each cluster combination to construct the contingency table and tested their association using a chi-square test of independence (chisq.test in R).

For the post-hoc analyses of each cell, we used adjusted Pearson residuals [26]to account for the unbalanced counts among the rows and columns in the contingency table for inter-brain clusters and behavioral clusters. Adjusted Pearson residuals are asymptotically normal, unlike ordinary Pearson residuals, which can underestimate sample variability and are not always normal in real data [10]. This supports more accurate inference at the cell level. They also serve as standardized effect sizes, with larger absolute values indicating stronger deviations from expected counts (positive means more observations than expected, negative means fewer). The formula is given in Supplementary Methods S4.

Cramér’s V was computed as the effect size, with a 95% confidence interval from bootstrap resampling of epochs (Supplementary Methods S4).

The unit of observation is the epoch. Epochs are nested within trials within dyads, which the Spearman and chi-square tests treat as independent, inflating the effective N and potentially deflating *p*-values. We therefore repeated the Spearman analyses at coarser aggregation and against a within-trial circular-shift null (Supplementary Methods S3, S7 Fig), and re-evaluated the cluster association with a mixed-effects logistic regression and a within-trial circular-shift permutation test (Supplementary Methods S4).

All *p* values were corrected for multiple comparisons with the Benjamini-Hochberg procedure, implemented via the fdr_bh function in MATLAB. This controlled the false discovery rate at 0.05.

### Software and Hardware

The EEG pre-processing and IBS computation were run in Python 3.11. The chi-square test was run in R 3.6.3. The mixed-effects logistic regression and the permutation null were run in MATLAB R2022b. The rest of the analyses were run in MATLAB 2022b (MathWorks Inc., Natick, MA, USA). The EEG pre-processing, IBS computation and dyad-shuffled surrogate method were run on the computing clusters at OIST.

## Results

### Analysis of Behavioral Patterns

We applied k-means clustering to the seven behavioral variables (S2 Fig) to characterize behavioral dynamics during the PCE. Fig 2A shows the normalized features defining the four clusters, and Fig 2B their within-trial dynamics at 3 s resolution. Clusters 1 and 2 represented the close-together modes, with the smallest inter-avatar distances (.136 and .154, respectively) and the highest mutual contact (.457 and .398, respectively). They differed mainly in the distance between each player and the partner’s shadow object (.955 and .203, respectively). Clusters 3 and 4 represented the far-apart modes. Cluster 3 had an intermediate inter-avatar distance (.501) with almost no mutual contact (.050), whereas cluster 4 had the largest inter-avatar distance (.647) and the smallest distance to the own static object of the participants (.271). Velocity differences were highest in cluster 3 (.190) and similar across the other clusters (.112, .139, and .134). Acceleration differences were highest in cluster 2 (.058) and lowest in cluster 1 (.028). DTW distances were broadly comparable across all four clusters (.214, .209, .190, and .234).

**Fig 2.**
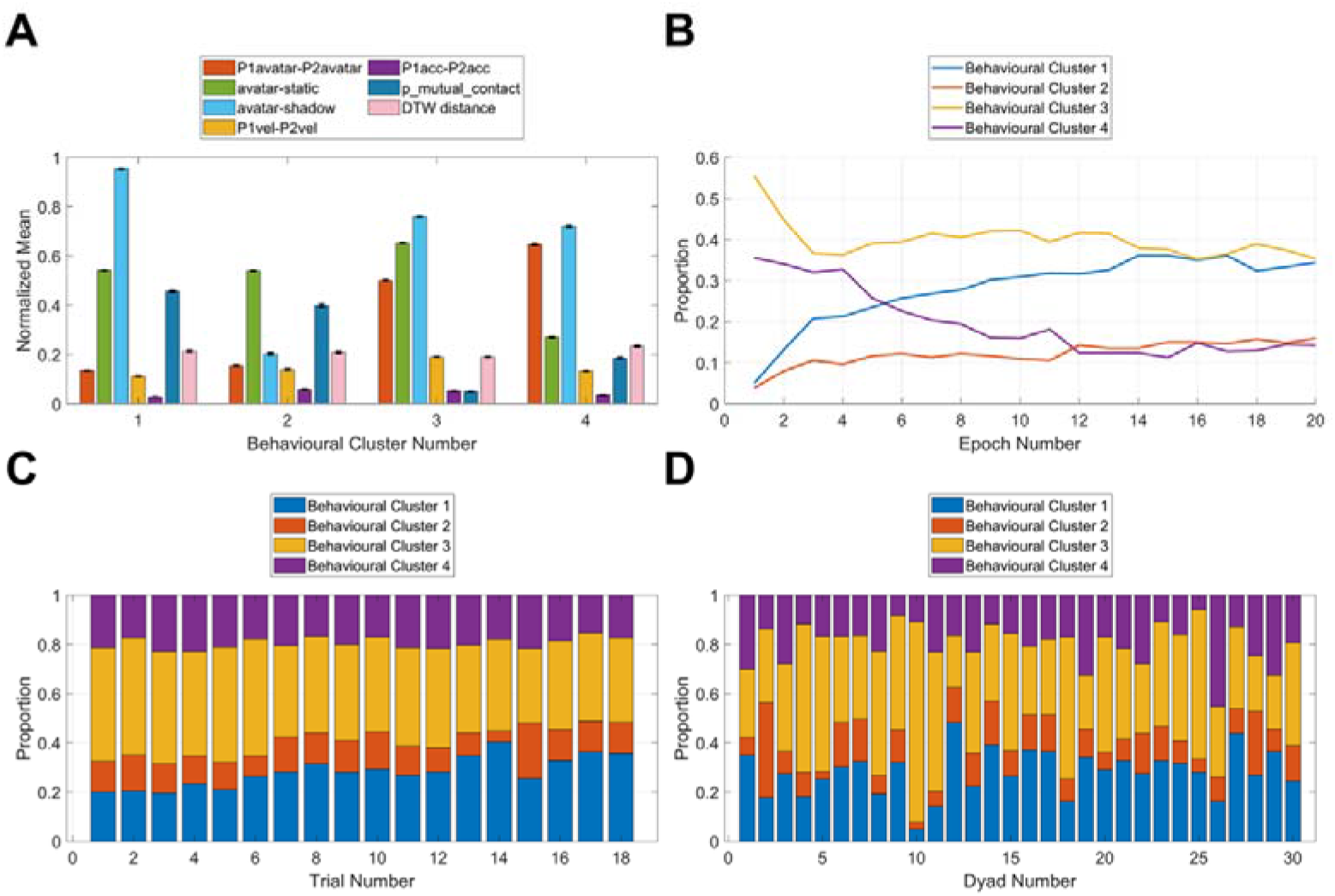
Behavioral pattern clusters identified using k-means and their dynamics. (A) Characteristics of each identified cluster. Each variable were averaged across all 3 s-epochs and normalized to the range [0,1]. avatar, participant avatar position; static, participant static object position; shadow, participant shadow object position; vel, velocity; acc, acceleration; DTW, dynamic time warping. Differences were computed as absolute differences. (B) Proportion for each cluster within a trial. Each epoch was 3 s. Clusters were color-coded for B, C and D. (C) Proportion for each trial across all dyads and epochs. (D) Proportion for each dyad across all trials and epochs.

The temporal dynamics in Fig 2B show that clusters 1 and 2 both increased in occurrence over the course of each trial, with cluster 1 increasing from .050 to approximately .343 and cluster 2 from .039 to approximately .152. Most of the change occurred within the first five epochs (0–15 s), after which the proportions changed more gradually. Cluster 2 reached a relatively stable level by approximately epoch 12 (33–36 s), and cluster 1 by approximately epoch 13 (36–39 s). Cluster 1 increased progressively to become one of the two dominant clusters in the later epochs, reaching a proportion comparable to that of cluster 3 (.343 vs. .367).

Clusters 3 and 4 showed the opposite pattern. Both had relatively high proportions at the onset of each trial (.556 and .356, respectively), which decreased over time to approximately .367 and .139, respectively. Cluster 3 reached a relatively stable level earlier in the trial (epoch 3, 6–9 s) than cluster 4 (epoch 10, 27–30 s). By the end of the trial, cluster 4 had the lowest proportion of the four clusters (.139), followed by cluster 2 (.152).

When averaged across the whole trial, the ordering differed because cluster 4 began at a relatively high proportion, whereas cluster 2 began at a relatively low proportion. Fig 2C shows that cluster proportions were largely independent of trial number, whereas interaction patterns differed substantially across dyads (Fig 2D). Across dyads, cluster 3 had the highest mean proportion (.3998 ± .0536), followed by cluster 1 (.2824 ± .0859), cluster 4 (.1953 ± .0822), and cluster 2 (.1225 ± .0292).

### Inter-brain Synchronization in the Alpha-Low and Alpha-High Bands

We computed CCorr for each 3 s epoch between the EEG sensor signals of the 30 dyads. To separate inter-brain coupling from similarity driven by the shared task structure, the observed values were ranked against a dyad-shuffled surrogate distribution and thresholded. A sensor pair reached the 97.5th percentile at 22 of the 871 possible ranks, corresponding to a nominal chance rate of 22/871 = .0253. Binary surrogate-thresholded connections were summed across epochs into density maps, and weak connections were removed using a percentile filter. This filter thresholds observed densities rather than testing significance, so retained connections are the strongest in the observed set, not individually statistically significant. Across sensor pairs and frequency bands, the observed proportions of surrogate-thresholded epochs ranged from .0247 to .0345, with a grand mean of .0266.

Figure 3 shows the strongest inter-brain connections after percentile filtering. In Figure 3A, which displays the top 0.5% of connections, the majority of inter-brain connections were observed in the Alpha-Low and Alpha-High frequency bands. Notably, the connections involved the frontal, central, temporal and parietal regions. Next, filtering at 0.1% gave a clearer picture of the connections (Fig 3B). There were more temporal connections in the Alpha-High band than the Alpha-Low band: at the sensor level the two bands differ in which pairs are strongest. Eight of the fifteen strongest Alpha-Low pairs involve Pz, P1 or P2 paired with frontal sensors, whereas eleven of the fifteen strongest Alpha-High pairs involve T8 or TP8 paired with left frontal or left temporal sensors. Because the PCE has no directionality (neither participant was instructed to lead), we averaged symmetrical connections across the upper and lower triangles of the matrices, and averaged sensor connections into ROIs shown as connection densities (Fig 3C). For Alpha-Low, the retained ROI-ROI connections were MID-MID (.031), RF-MID (.027), RC-MID (.013), RF-LP (.009), LF-LP (.009), RT-LT (.008), RP-MID (.008) and LP-MID (.008). For Alpha-High they were RT-LF (.045), RT-LT (.031), RF-RT (.018), RF-LP (.009), RF-MID (.009) and LF-MID (.009). Alpha-Low was therefore dominated by midline connections and Alpha-High by right temporal connections.

**Fig 3.**
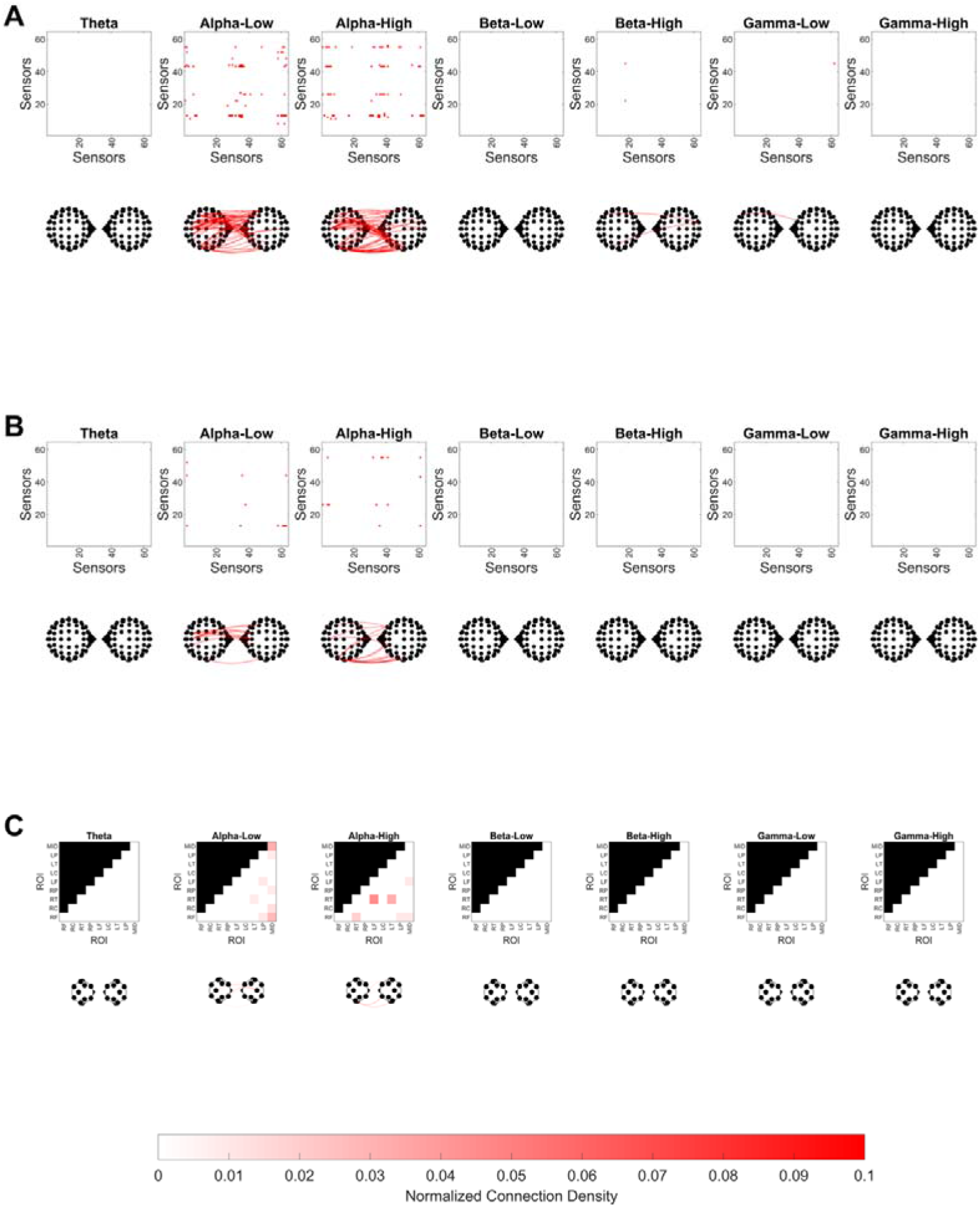
Inter-brain connections retained by the dyad-shuffled surrogate method. The surrogate test is applied per epoch, per frequency band and per sensor pair. (A) Connections in the top 0.5% of all sensor pairs taken across the seven frequency bands together. Each red dot on a matrix marks a retained sensor-sensor connection, and each red curve on a head plot marks that connection between the two heads, which face each other. The bands are Theta (4 to 7.5 Hz), Alpha-Low (7.5 to 10 Hz), Alpha-High (10 to 12.5 Hz), Beta-Low (12.5 to 20 Hz), Beta-High (20 to 30 Hz), Gamma-Low (30 to 58 Hz) and Gamma-High (62 to 100 Hz). (B) The same, retaining the top 0.1%. (C) The top 0.1% sensor-sensor connections averaged into nine regions of interest (ROI-ROI) connections. Symmetrical connections were averaged. The color scale shows the number of connections normalized by the total number of connections within each ROI-ROI. Because ROI blocks differ in size, from 25 to 64 sensor pairs, an equal fraction corresponds to a different number of retained connections in each block, and cells in smaller blocks reach high values with fewer connections.

To test whether these ROI connections related to behavior, we computed Spearman correlations between each retained connection (eight for Alpha-Low, six for Alpha-High) and the seven behavioral variables (Table 1). After FDR correction across all connections and behavioral variables, 27 cells survived, 14 in Alpha-Low and 13 in Alpha-High. Inter-avatar distance was positively correlated with every retained connection in both bands, with *ρ* ranging from .033 to .049 in Alpha-Low and from .027 to .038 in Alpha-High. The proportion of mutual contact was negatively correlated with four of the eight Alpha-Low connections and three of the six Alpha-High connections, with *ρ* ranging from -.030 to -.024. In Alpha-Low, avatar–static-object distance and avatar–shadow distance were each negatively correlated with one connection. In Alpha-High, velocity difference was negatively correlated with four of the six connections. All correlation coefficients were very small. Because epochs are not independent observations, we additionally evaluated each correlation against a within-trial circular-shift null distribution (Supplementary Methods S3). Under this null, none of the 27 previously significant cells survived multiple-comparison correction (smallest adjusted permutation *p* = .28), indicating that the apparent epoch-level associations were not robust once the temporal dependence structure of the data was taken into account.

**Table 1.**
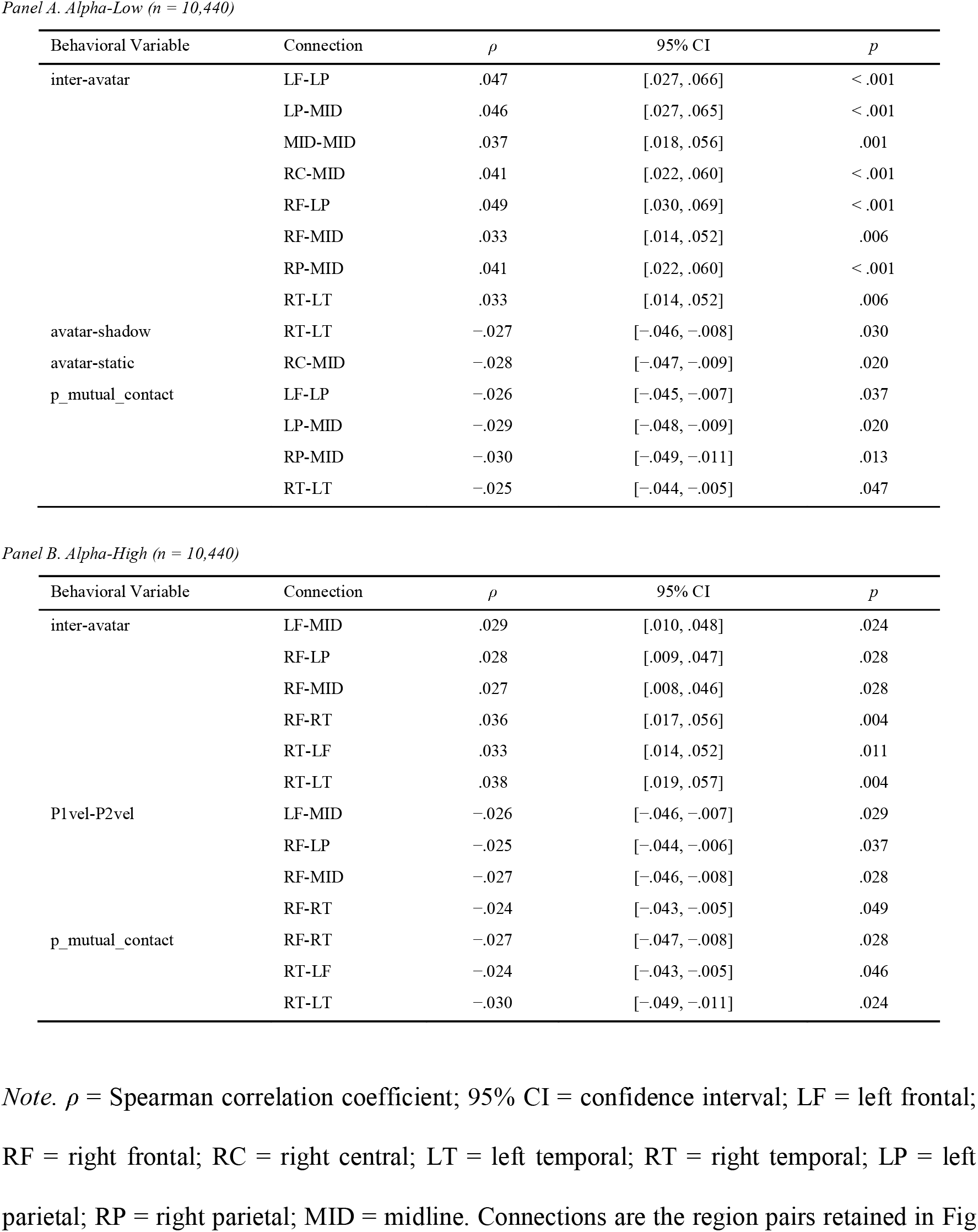
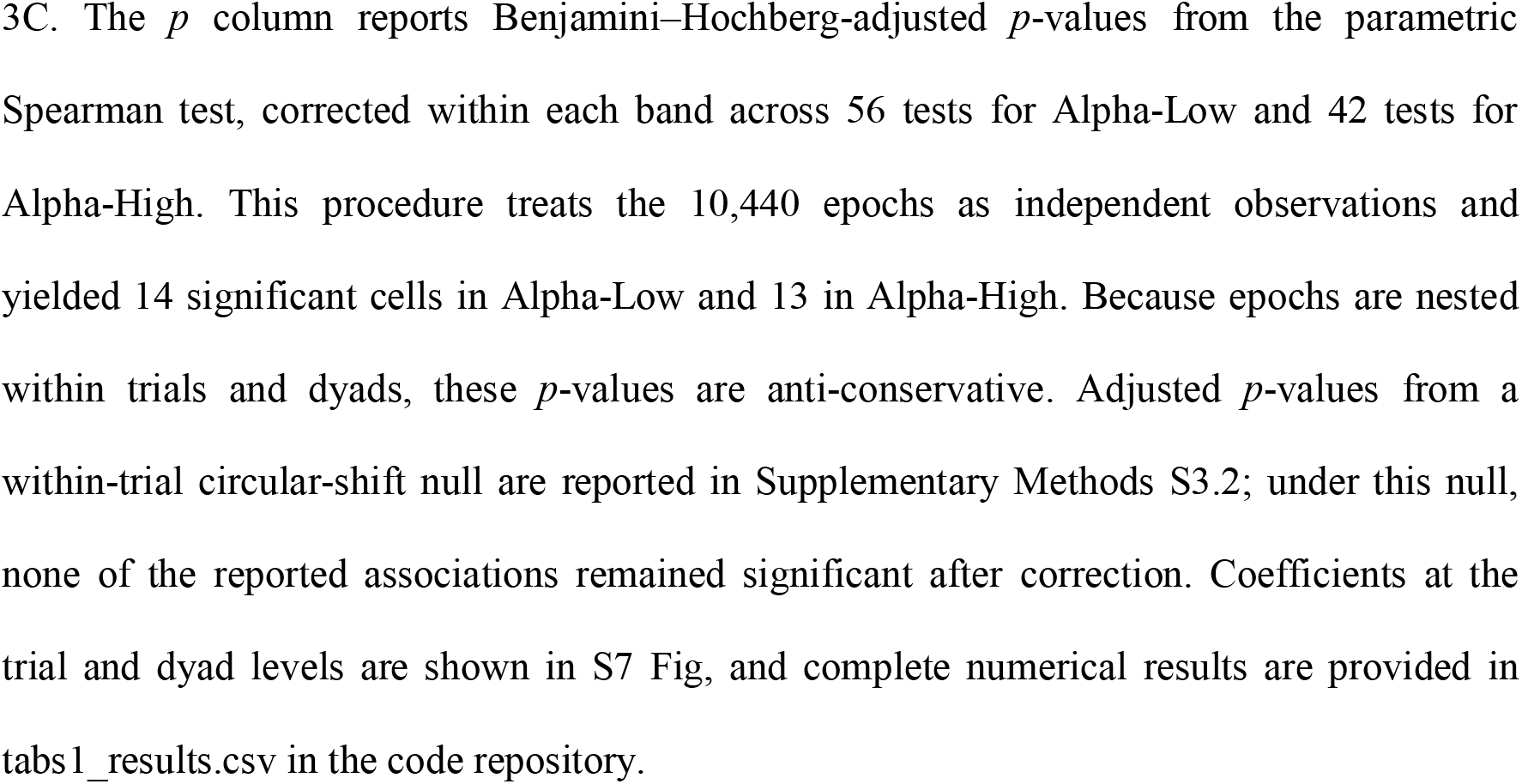
Spearman Correlations Between Behavioral Variables and Inter-brain Connections Surviving Correction in Alpha Bands.

### Inter-brain Connectivity Patterns

Similarly to the behavioral analysis, for the Alpha-Low and Alpha-High frequency bands, we applied k-means clustering to the full inter-brain FC thresholded using the surrogate method. The optimal number of clusters was found to be three. After labeling each epoch with a cluster and averaging connectivities per cluster, we kept only the top 0.5% of connections per cluster. The cluster maps therefore use the top 0.5%, whereas the pooled map in Fig 3C uses the top 0.1%, so the two are not directly comparable. Retained connections were averaged into the nine ROIs and symmetrized across participants, since which member is P1 is arbitrary. Fig 4 and Fig 5 shows the resulting connectivity patterns for each inter-brain cluster and their dynamics. Fig 4A shows the connectivity for each inter-brain cluster together with an all-epoch reference block, and the three clusters had different spatial configurations. Cluster 1 was by far the most prevalent (9,541 of 10,440 epochs, 91.4%) and its connections were concentrated in the temporal regions, both within each hemisphere (LT-LT, .078; RT-RT, .047) and between them (RT-LT, .039). Cluster 2 (699 epochs, 6.7%) was the most diffuse: a left fronto-parietal connection was strongest (LF-LP, .036) and its remaining ten connections were spread across frontal, temporal, parietal and midline regions with no value above .018. Cluster 3 was the rarest (200 epochs, 1.9%) and was confined to the frontal and midline regions (LF-MID, .054; MID-MID, .047; RF-RF, .041).

**Fig 4.**
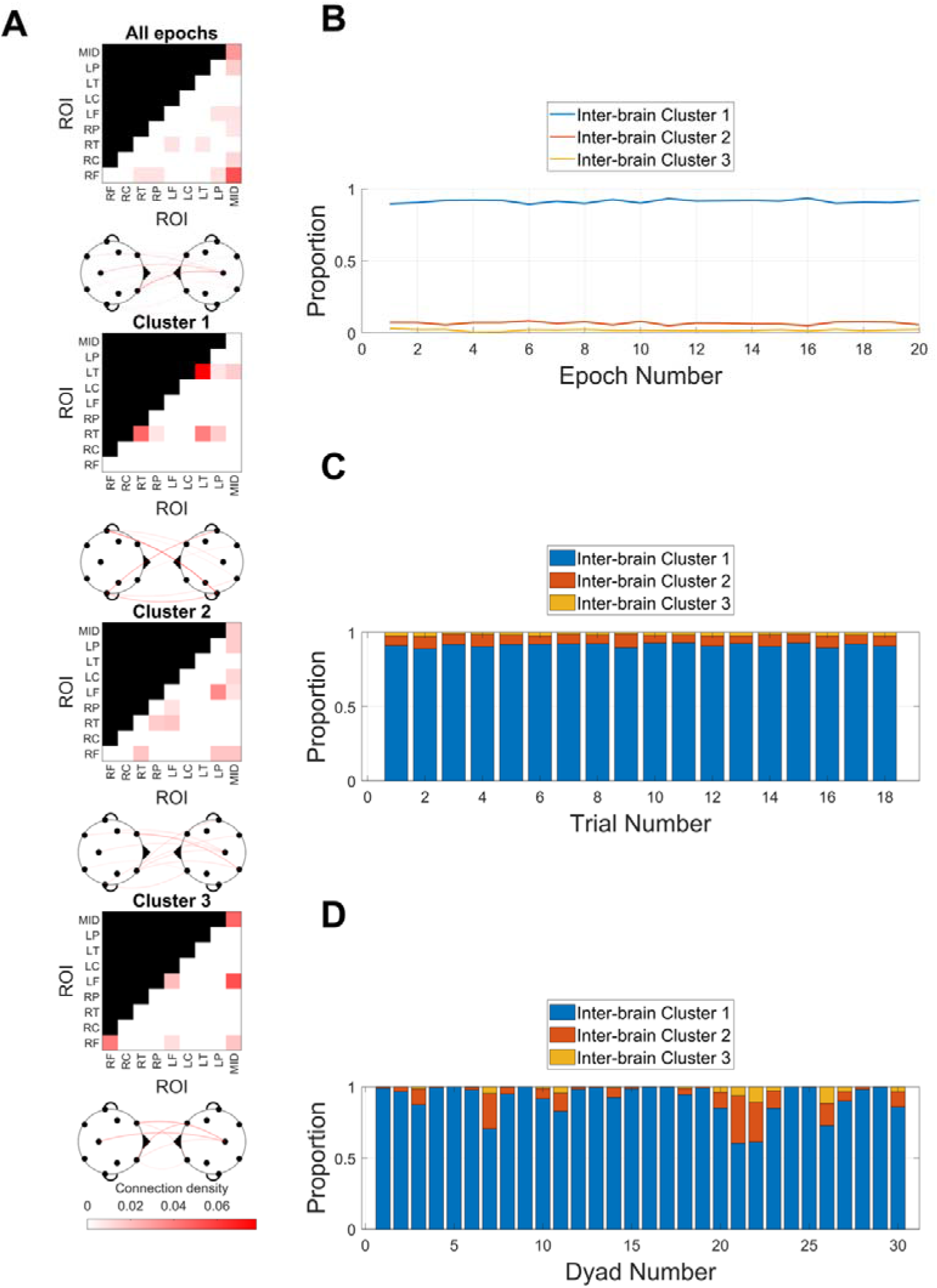
Inter-brain connectivity pattern clusters identified using k-means and their dynamics for the Alpha-Low band. (A) Inter-brain clusters and connectivity for the Alpha-Low (7.5-10 Hz) frequency band. The top block serves as a reference for all epochs and the color scale is shared by all four blocks. After summing up connectivity patterns across all epochs corresponding to each cluster, sensor-sensor connectivity patterns were obtained by retaining the top 0.5% of connections within each cluster, and were averaged into nine regions of interest (ROI-ROI). (B) Proportion of each cluster over epochs within a trial. Each epoch was 3 s. Clusters were color-coded for B, C and D. (C) Proportion for each trial across all dyads and epochs. (D) Proportion for each dyad across all trials and epochs.

**Fig 5.**
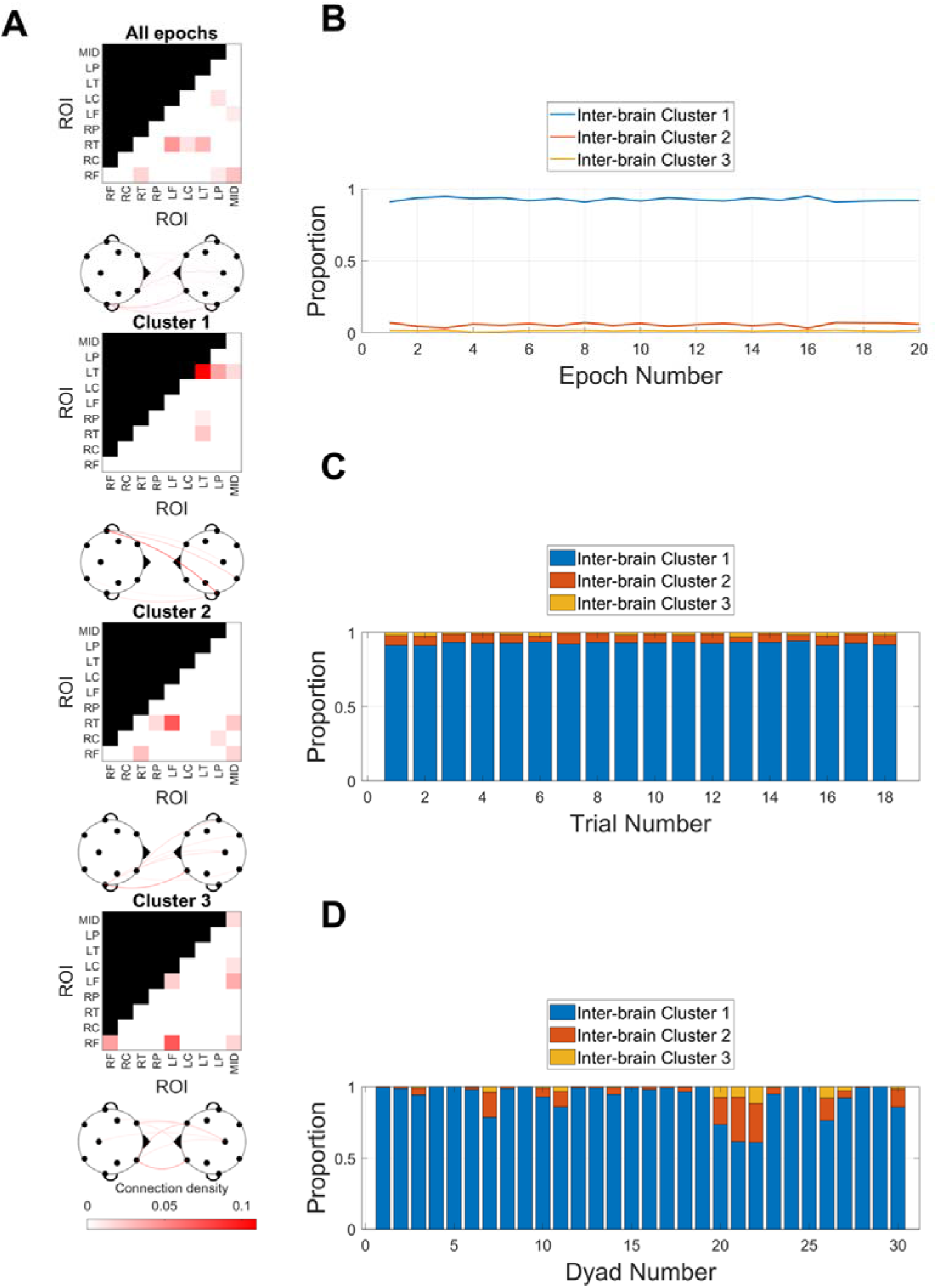
Inter-brain connectivity pattern clusters identified using k-means and their dynamics for the Alpha-High band. (A) Inter-brain clusters and connectivity for the Alpha-High (10.0-12.5 Hz) frequency band. The top block serves as a reference for all epochs and the color scale is shared by all four blocks. After summing up connectivity patterns across all epochs corresponding to each cluster, sensor-sensor connectivity patterns were obtained by retaining the top 0.5% of connections within each cluster, and were averaged into nine regions of interest (ROI-ROI). (B) Proportion of each cluster over epochs within a trial. Each epoch was 3 s. Clusters were color-coded for B, C and D. (C) Proportion for each trial across all dyads and epochs. (D) Proportion for each dyad across all trials and epochs.

The proportions of these three clusters were consistent within and across trials (Fig 4B, C) but varied across dyads (Fig 4D, Fig 5D), especially for clusters 2 and 3, as with the behavioral clusters. In Alpha-Low the between-dyad *SD*s were .115, .088 and .031 against means of .914, .067 and .019. Both minority clusters had *SD*s larger than their own means. Cluster 2 ranged from .000 to .332 across dyads and cluster 3 from .000 to .117. In Alpha-Low, 3 of 30 dyads never entered cluster 2 and 12 of 30 never entered cluster 3, yet cluster 2 accounted for as much as .332 of the epochs of a single dyad. Alpha-High showed the same pattern, with 7 and 15 dyads respectively.

An exploratory analysis was conducted to examine the relationship between the JS rate and dyadic proportion in each cluster, but no significant correlations were found in either band. For Alpha-Low, cluster 1, *ρ* = .0965; cluster 2, *ρ* = -.0918; cluster 3, *ρ* = -.1780; all adjusted *p* = .6295. For Alpha-High, cluster 1, *ρ* = .0901; cluster 2, *ρ* = -.0905; cluster 3, *ρ* = - .0342; all adjusted *p* = .8576.

Alpha-High patterns and dynamics broadly matched those of the Alpha-Low band (Fig 5), with the same ordering of cluster prevalence (9,677, 593 and 170 epochs) and the same three-way distinction between a dominant temporal cluster, an intermediate cluster involving frontal and temporal regions, and a rare frontal-midline cluster. Cluster 1 was almost exclusively left temporal in Alpha-High (LT-LT, .109; LT-LP, .039), whereas in Alpha-Low it involved both hemispheres (LT-LT, RT-RT and RT-LT). Cluster 2 was a right temporal to left frontal connection in Alpha-High (RT-LF, .071) but left fronto-parietal and considerably more diffuse in Alpha-Low (LF-LP, .036).

### Exploration of the Association Between Behavioral and Inter-brain Clusters

The frequency of occurrence for each inter-brain and behavioral cluster is reported in Table 2. To assess how the two cluster types related, we computed conditional proportions in both directions: each behavioral cluster given each inter-brain cluster, and each inter-brain cluster given each behavioral cluster. Visual inspection of the conditional proportions suggested that inter-brain cluster 1 dominated across all four behavioral clusters, at .925, .925, .910 and .899 for behavioral clusters 1 to 4 in Alpha-Low, so the inter-brain distribution is close to flat when conditioned this way (Fig 6A). Conditioned on the inter-brain cluster (Fig 6B), the behavioral composition shifted monotonically. From inter-brain cluster 1 to 3, behavioral cluster 1 became less frequent (.286, .255, .215) and behavioral cluster 2 also declined (.125, .110, .100), while behavioral cluster 3 (.398, .409, .445) and behavioral cluster 4 (.191, .226, .240) became more frequent. Alpha-High is nearly identical (.934, .942, .923, .914 for Fig 6C; and .285 to .235, .398 to .429 for Fig 6D). Because behavioral cluster 1 is the closest-together, highest mutual-contact mode while clusters 3 and 4 are the far-apart modes, the two less common inter-brain clusters tended to occur when the participants were further apart and in less mutual contact.

**Fig 6.**
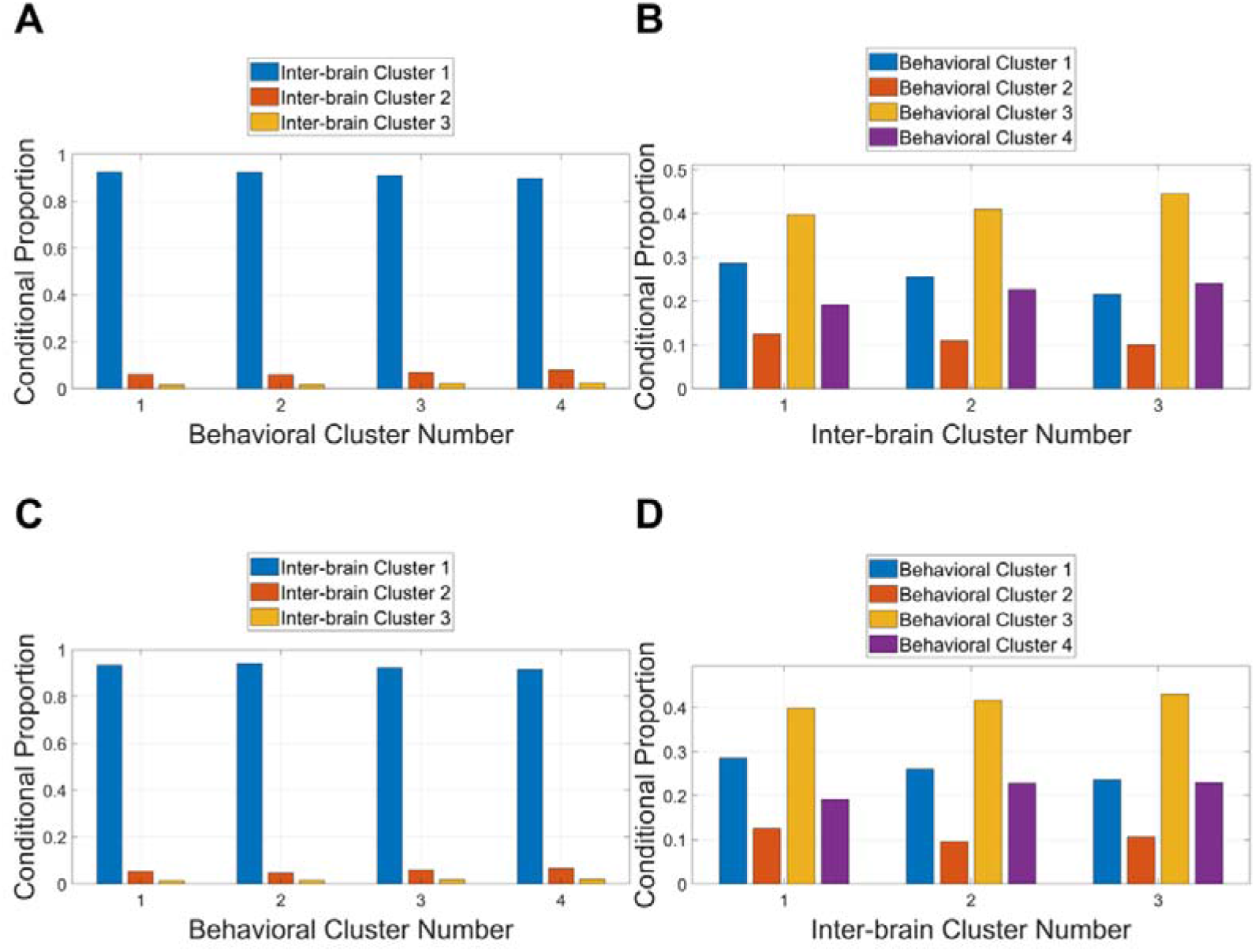
Conditional proportions of behavioral and inter-brain clusters. (A, C) Proportion of inter-brain clusters conditioned on behavioral clusters for the Alpha-Low (A) and Alpha-High (C) bands. (B, D) Proportion of behavioral clusters conditioned on inter-brain clusters for the Alpha-Low (B) and Alpha-High (D) bands. Proportions were computed across all epochs, trials, and dyads.

**Table 2.** Frequencies of Behavioral and Inter-brain Cluster Combinations by Alpha Band.

| Behavioral<br>Cluster | Inter-brain Cluster |  |  |  |  |  |  |  |
| --- | --- | --- | --- | --- | --- | --- | --- | --- |
|  | Alpha-Low |  |  |  | Alpha-High |  |  |  |
|  | 1 | 2 | 3 | Total | 1 | 2 | 3 | Total |
| 1 | 2732 | 178 | 43 | 2953 | 2759 | 154 | 40 | 2953 |
| 2 | 1191 | 77 | 20 | 1288 | 1213 | 57 | 18 | 1288 |
| 3 | 3793 | 286 | 89 | 4168 | 3848 | 247 | 73 | 4168 |
| 4 | 1825 | 158 | 48 | 2031 | 1857 | 135 | 39 | 2031 |
| **Total** | 9541 | 699 | 200 | 10440 | 9677 | 593 | 170 | 10440 |

*Note.* Frequencies represent the number of occurrences for each behavioral and inter-brain cluster combination under Alpha-Low and Alpha-High conditions.

To quantify the association between behavioral and inter-brain clusters, we applied a chi-square test of independence to Table 2, which was significant (Alpha-Low, □^2^(6) = 15.17, *n* = 10,440, *p* = .019; Alpha-High, □^2^(6) = 12.60, *n* = 10,440, *p* = .0498), neither of which supported the association once the nesting of epochs was taken into account (see below). Cramér’s V indicated very small effect sizes for both frequency bands (Alpha-Low V = .027, 95% CI [.019, .044]; Alpha-High V = .025, 95% CI [.017, .041]).

Post-hoc local analyses based on adjusted Pearson residuals (Table 3; Table 4) revealed no significant associations between any of the behavioral clusters and inter-brain clusters. The largest adjusted residual was observed for the pairing of behavioral cluster 4 and inter-brain cluster 1, and it was negative (Alpha-Low: residual = −2.74, *p* = .060; Alpha-High: residual = −2.43, *p* = .113), indicating a tendency for the far-apart behavioral mode to co-occur less often than expected with the dominant inter-brain cluster. No single cell drives the omnibus association: any departure from independence is small and spread across the table.

**Table 3.** Post-Hoc Adjusted Pearson Residuals from the Chi-Square Test of Independence Between Behavioral and Inter-brain Clusters in the Alpha-Low Band, with Theoretical and Permutation p-values (n = 10,440)

| Behavioral Cluster | Inter-brain Cluster 1 | Inter-brain Cluster 2 | Inter-brain Cluster 3 |
| --- | --- | --- | --- |
| 1 | 2.58 ($p = .060$;<br>$p_{\text{perm}} = .255$) | -1.71 ($p = .202$;<br>$p_{\text{perm}} = .405$) | -2.15 ($p = .094$;<br>$p_{\text{perm}} = .255$) |
| 2 | 1.48 ($p = .240$;<br>$p_{\text{perm}} = .405$) | -1.10 ($p = .326$;<br>$p_{\text{perm}} = .405$) | -1.01 ($p = .338$;<br>$p_{\text{perm}} = .547$) |
| 3 | -1.15 ($p = .326$;<br>$p_{\text{perm}} = .442$) | 0.55 ($p = .579$;<br>$p_{\text{perm}} = .601$) | 1.33 ($p = .273$;<br>$p_{\text{perm}} = .405$) |
| 4 | -2.74 ($p = .060$;<br>$p_{\text{perm}} = .547$) | 2.18 ($p = .094$;<br>$p_{\text{perm}} = .448$) | 1.64 ($p = .202$;<br>$p_{\text{perm}} = .703$) |

*Note.* Values represent adjusted Pearson residuals and Benjamini-Hochberg-corrected $p$ -values from the theoretical chi-square reference distribution ( $p$ ) and the within-trial circular-shift null ( $p_{\text{perm}}$ ; 10,000 permutations; Supplementary Methods S4.2). No cell reached the .05 level after correction under either reference distribution.

**Table 4.**
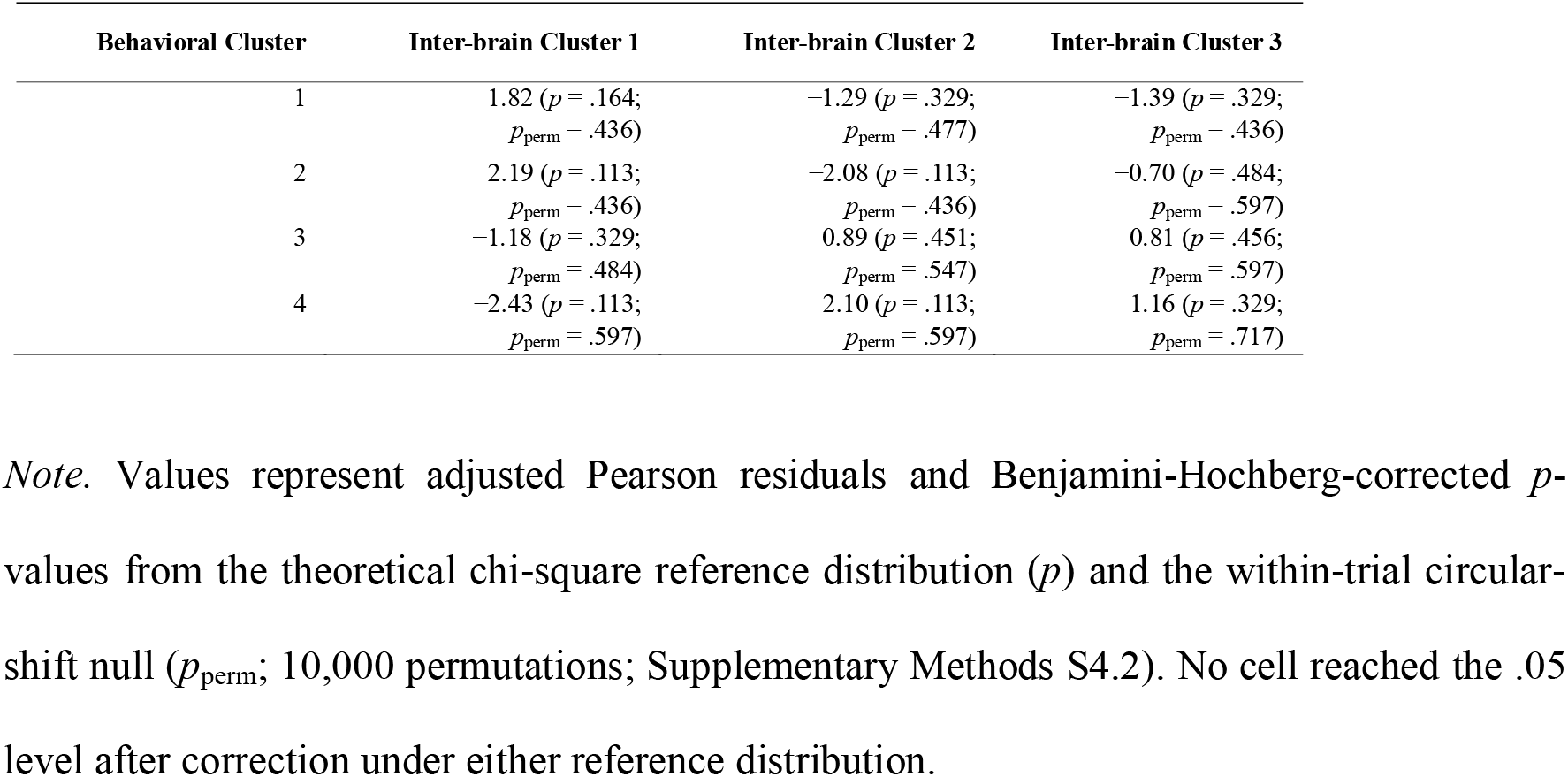
Post-Hoc Adjusted Pearson Residuals from the Chi-Square Test of Independence Between Behavioral and Inter-brain Clusters in the Alpha-High Band, with Theoretical and Permutation p-values (n = 10,440)

| Behavioral Cluster | Inter-brain Cluster 1 | Inter-brain Cluster 2 | Inter-brain Cluster 3 |
| --- | --- | --- | --- |
| 1 | 1.82 ($p = .164$;<br>$p_{\text{perm}} = .436$) | -1.29 ($p = .329$;<br>$p_{\text{perm}} = .477$) | -1.39 ($p = .329$;<br>$p_{\text{perm}} = .436$) |
| 2 | 2.19 ($p = .113$;<br>$p_{\text{perm}} = .436$) | -2.08 ($p = .113$;<br>$p_{\text{perm}} = .436$) | -0.70 ($p = .484$;<br>$p_{\text{perm}} = .597$) |
| 3 | -1.18 ($p = .329$;<br>$p_{\text{perm}} = .484$) | 0.89 ($p = .451$;<br>$p_{\text{perm}} = .547$) | 0.81 ($p = .456$;<br>$p_{\text{perm}} = .597$) |
| 4 | -2.43 ($p = .113$;<br>$p_{\text{perm}} = .597$) | 2.10 ($p = .113$;<br>$p_{\text{perm}} = .597$) | 1.16 ($p = .329$;<br>$p_{\text{perm}} = .717$) |

*Note.* Values represent adjusted Pearson residuals and Benjamini-Hochberg-corrected $p$ -values from the theoretical chi-square reference distribution ( $p$ ) and the within-trial circular-shift null ( $p_{\text{perm}}$ ; 10,000 permutations; Supplementary Methods S4.2). No cell reached the .05 level after correction under either reference distribution.

Because epochs are nested within trials within dyads, we repeated this test as a mixed-effects logistic regression predicting whether an epoch fell in one of the two less common inter-brain clusters from its behavioral cluster, with random intercepts for dyad and for trial within dyad; the behavioral effect was not significant in either band (Alpha-Low, □^2^(3) = 7.65, *p* = .054; Alpha-High, □^2^(3) = 6.90, *p* = .075; see Supplementary Methods S4). We also built an empirical null for the chi-square itself by circularly shifting the behavioral labels within each trial, which preserves the composition of every dyad and trial while breaking the alignment between the two label streams, and against this null the association was clearly not significant (Alpha-Low *p* = .307, Alpha-High *p* = .433; see Supplementary Methods S4).

## Discussion

To explore whether inter-brain coupling can occur during minimal tactile interaction between physically isolated individuals, we conducted an exploratory analysis of a publicly available PCE dataset. This work is descriptive and preliminary: it accounts for how behavioral and inter-brain dynamics may co-occur in a minimal interaction context rather than providing confirmatory evidence of specific neural mechanisms.

### Behavioral Modes in the PCE

Real-time coordination is best examined in paradigms that are active, continuous, and dynamic [55], yet sufficiently constrained for systematic analysis [38], criteria that the PCE satisfies. Because no single variable can capture the complexity of interaction, we computed multiple behavioral measures and used k-means clustering to identify recurring patterns in both behavior and inter-brain functional connectivity (FC). This approach is commonly used to classify brain states in the FC literature [3] and has more recently been applied in hyperscanning [8, 46].

Across analyses we identified four behavioral patterns (Fig 2A) and three inter-brain patterns in the alpha band (Fig 4A, Fig 5A). The proportion of time in each behavioral cluster was stable across trials (Fig 2C, Fig 4C) but differed across dyads (Fig 2D, Fig 4D). Within a trial, clusters 3 and 4 became less frequent while clusters 1 and 2 became more frequent. Cluster 3 fell rapidly in the first few epochs and then stabilized, whereas clusters 1, 2, and 4 changed more gradually. These patterns suggest an initial transient period of rapid change followed by a more stable distribution, and let us group the behavioral patterns into periods when participants were spatially closer (clusters 1 and 2) and periods when they were farther apart (clusters 3 and 4).

Notably, cluster 3 had the highest overall prevalence and was characterized by the lowest mutual contact and moderate inter-avatar distance. Rather than a clear strategy, it appears to be a weakly coupled mode reflecting the minimal, ambiguous sensory information in the PCE. It likely includes the starting configuration of each trial, but its persistence suggests an ongoing default mode rather than a transient state, from which more strongly coupled patterns (clusters 1 and 2) emerge as coordination develops. Clusters 1 and 2 involve close mutual engagement, with the smallest inter-avatar distances and highest mutual contact, and predominated later in each trial, whereas clusters 3 and 4 centered on objects other than the partner and were more prevalent early on. Cluster 4 is distinct: despite the largest inter-avatar distance, its mutual contact was elevated relative to cluster 3, suggesting intermittent avatar-avatar contact, and it had the smallest distance to the own static object.

### Alpha-band Inter-brain Connections and their Topography

Interestingly, we did not find a clear temporal progression in the inter-brain network patterns analogous to the behavioral convergence. Across a trial, the proportions of the three alpha-band inter-brain clusters remained broadly stable (Fig 4B; Fig 5B). This stability should be read with the caveat that one cluster accounted for most epochs in both bands. Descriptively, inter-brain configuration did not track the time-evolving behavioral coordination at the resolution examined. We did not test this correspondence formally and report it as an observation at the scale of a few seconds.

The connections retained by the percentile filter are the strongest in the observed set rather than individually significant ones, so they are compatible with, but do not establish, phase-locked activity between partners who have no visual contact, are physically separated, and share only a minimal tactile interaction. The alpha bands were the only ranges in which any connection survived the percentile filter (top 0.1%, Fig 3B). Alpha synchronization has been reported in other physically isolated setups, including alpha-mu (8-12 Hz) coupling during imitation in separate rooms [16] and alpha coupling in a collaborative online game [69], and IBS has been reported in remote video interactions, such as frontal-temporal beta synchrony between separated individuals [64]. Across tasks, IBC is commonly reported in theta, alpha, and beta, with some gamma reports [4, 35, 36, 43, 54, 61, 67, 70], though EEG gamma hypersynchrony is artifact-prone [30].

The strongest inter-brain connections differed in their spatial distribution across the two alpha sub-bands: retained Alpha-Low connections were concentrated on the midline, while retained Alpha-High connections were primarily over the right temporal region (Fig 3B, Fig 3C). These patterns are descriptive, as the percentile filter identifies the strongest observed connections rather than individually significant ones.

The Alpha-High pattern is broadly consistent with reports of temporo-parietal inter-brain coupling during social interaction [13, 66], though both used fNIRS, and the same meta-analysis reported prefrontal coupling at least as consistently [13]. Regions represented here, including the rTPJ, have been linked to self-other distinction and social cognition [1]. The right-lateralized pattern is also consistent with the tendency for hyperscanning studies to report right-hemisphere coupling [64] and with evidence for a right-sided 10–12 Hz mechanism in intentional social coordination, although this evidence concerns within-brain rather than inter-brain synchronization [55]. Posterior-frontal coupling in this frequency range has also been reported, with right parietal–prefrontal connectivity modulated by social judgment [15].

The Alpha-Low pattern has less direct precedent. Low-alpha IBS has been reported in EEG hyperscanning during social imitation [52], as has alpha coupling over parietal sensors, including CPz, Pz, and adjacent sites, during a cooperative online game involving spatially separated participants [69]. Another relevant comparison is alpha-mu coupling over right centro-parietal sensors during spontaneous imitation [16]. However, that pattern was right-lateralized, whereas our Alpha-Low pattern was centered on the midline.

Probabilistic mappings of scalp electrodes to brain regions [65] indicate that several RT sensors overlie parts of the inferior, middle, and superior temporal gyri and the angular gyrus. However, the RT region also includes more anterior, central, and posterior sites, and these correspondences vary across individuals, particularly for central and parietal electrodes [65]. These temporal and parietal structures have been linked to processes such as recognition, motion perception, auditory processing, multisensory integration, and attentional orienting [28]. Similar areas have been linked to mirror-neuron and motor-resonance networks for social perception and coordination [55], though our task lacked the visual or action cues that typically engage such circuitry. Given the absence of source localization and the limited spatial precision of EEG, any correspondence with specific neural systems remains suggestive rather than definitive.

Taken together, these scalp-level patterns are compatible with shared task engagement or attentional alignment rather than activation of a specific cortical system, with the Alpha-High pattern overlapping regions implicated in social-attentional processing. We next asked whether the two alpha sub-bands differed in their connections, since low and high alpha have been proposed to reflect distinct processes.

The framing that motivated this study, in which social touch is associated with alpha coupling over somatosensory regions, does not describe what we observed. The retained connections lay along the midline and over the right temporal region rather than over frontal and central sensors, and descriptively higher coupling accompanied greater separation and less contact, without surviving the within-trial null (Supplementary Methods S3.2). The design cannot distinguish two interpretations. The tactile signal here is a discrete cue from a hand-held device rather than skin-to-skin contact, so it may not engage the same somatosensory processes as direct touch. Alternatively, the retained connections may reflect the shared search for one another rather than contact itself, so periods of greatest separation may also be periods of greater shared engagement.

The two sub-bands showed broadly similar overall topographies and differed mainly in their most prominent connections. Intra-brain and mu-suppression studies indicate that both low- and high-alpha contribute to sensorimotor and social processing [17], and others report differential alpha involvement during action observation and social interaction [20, 55]. We cite this to support the sub-band distinction, not as evidence for a mechanism in the PCE. At the inter-brain level, low-alpha has been reported as more prominent during demanding interactions and high-alpha during passive viewing in neurotypical dyads [52]. These findings do not support a one-to-one mapping between sub-bands and cognitive processes, and our differences were small. We therefore do not assign distinct functional roles to Alpha-Low and Alpha-High IBS but present the comparison descriptively. A functional interpretation would require a design that directly manipulates the processes of interest.

### Relating Behavior to Inter-brain Configurations

A key part of our analysis examined the relationship between the neural and behavioral patterns. The distribution of inter-brain configurations appeared to depend on the behavioral pattern in a conventional chi-square test. However, no specific pairing survived the post-hoc test, and this association did not survive either of two analyses that account for the nesting of epochs within trials and within dyads (Supplementary Methods S4). We therefore treat the association as descriptive and do not interpret specific pairings of behavioral and inter-brain clusters.

Prior work has linked frontal inter-brain coupling to social coordination, attentional regulation, and executive processes during joint action [13]. The present findings neither support nor exclude such a link. Future studies should separate inter-brain dynamics during interaction with the partner avatar from those with shadow objects, to test whether social context modulates frontal and temporal coupling.

Our correlation analyses between specific inter-brain connections and continuous behavioral measures further underscore the subtle nature of these links. The correlations were very small and nominally significant (Table 1), but none of the 27 associations survived the within-trial permutation null (minimum adjusted *p* = .28; Supplementary Methods S3.2). Descriptively, in both bands, inter-avatar distance was positively correlated with coupling strength on every retained connection, while the proportion of mutual contact was negatively correlated with coupling strength on several connections. Thus, interaction phases in which the partners were farther apart and in less direct contact were associated with stronger retained coupling. In the Alpha-High band, velocity difference was also negatively correlated with coupling strength on most connections, indicating that stronger coupling was associated with more similar movement speeds. We do not assign a functional interpretation to these associations because similar movement could produce similar alpha dynamics without interaction, and the design lacks a non-interactive condition to distinguish shared sensorimotor states from interaction-related coupling.

### Limitations

The observed association and correlation effect sizes were small, and their interpretation should consider the level of analysis. The PCE produces continuous interaction data with gradual transitions, temporal dependence, and substantial between-dyad variability. Epoch-level measures may therefore capture weak aggregate associations rather than strong or predictive relationships, while a 3-s epoch may combine multiple behavioral micro-states. Finer temporal analyses may better capture transient inter-brain dynamics [8].

Epochs were nested within trials and dyads, violating the independence assumption of the epoch-level Spearman and chi-square tests and inflating the nominal sample size. For the Spearman analyses, we addressed this in two ways (Supplementary Methods S3, S7 Fig): by repeating the analyses at coarser levels of aggregation and by evaluating epoch-level associations against a within-trial circular-shift null that preserves all epochs while accounting for temporal dependence. Correlation coefficients increased with aggregation, but this increase is expected as the number of observations decreases, so the aggregated analyses neither support nor rule out a dyad-level association. Under the within-trial null, no epoch-level association remained significant after correction, indicating that the associations were not robust to temporal dependence. For the cluster associations, trial-level aggregation was not feasible because one inter-brain cluster dominated both bands, leaving most trials assigned to a single cluster. We therefore used mixed-effects logistic regression with random intercepts for dyad and trial within dyad, together with a permutation null that circularly shifted behavioral labels within each trial (Supplementary Methods S4). Neither analysis supported the association. The nominal chi-square findings were therefore not robust to analyses accounting for the hierarchical structure, and the results in Tables 3 and 4 should be interpreted as descriptive rather than as evidence of reliable associations.

A further limitation concerns the relationship between IBS, task performance, and interpersonal attunement. We found no correlation between the JS metric, our primary measure of task performance, and the occurrence of any inter-brain cluster. This provides no evidence that IBS was directly related to explicit task success. Given the high dimensionality and noise of hyperscanning EEG, weak associations are not unexpected [11, 73]. The original dataset includes post-task ratings of interpersonal attunement, but these were not analyzed here, leaving open whether greater IBS was associated with greater subjective attunement. The absence of control conditions (solitary interaction, interaction with an artificial agent, or a non-social baseline) also limits whether the observed synchrony is specific to human-human interaction rather than general task engagement. The observed behavioral associations should therefore be interpreted as preliminary indicators of coordination rather than markers of social attunement. Future work should analyze the subjective data and incorporate appropriate control conditions to clarify the significance of IBC.

### Conceptual Implications

These implications extend beyond what the present data support. Behavioral coordination developed within trials even in an unfamiliar virtual environment with a single haptic channel, while inter-brain coupling remained weak and descriptive. That pattern lends itself to further interpretative work in the context of early embodied-cognitive-science views that virtual spaces cannot host authentic social interaction [22, 49], as well as more recent theoretical research that tends to recognize the constructive potential of virtual spaces for the emergence of new forms of mediated behavioral coordination [32, 33, 57, 58, 59, 68].

## Data Availability Statement

All scripts including scripts for generating the tables and figures, as well as links to the complete (raw, pre-processed and analyzed) dataset could be accessed freely at https://gitlab.com/oist-ecsu/ibspce2024.

## Acknowledgments

We are grateful for the computational resources provided by the Scientific Computing and Data Analysis section of Research Support Division at OIST.

## Declaration of Generative AI Use

The authors used generative AI tools for copyediting and language refinement, idea generation and exploration, and software coding assistance. The authors reviewed and verified all outputs and take full responsibility for the content of the manuscript.

## Supplementary Materials

### Supplementary Methods

#### S1. Behavioral clustering: optimal number of clusters and preprocessing

In the behavioral and inter-brain data analysis, we identified the optimal number of clusters using the elbow method. First, we plotted the elbow plot of clustering cost against the number of clusters (S3 Fig). Subsequently, the optimal number of clusters was determined automatically by locating the elbow of the plot. The elbow was defined as the point having the maximum perpendicular distance from the straight line between the first and the last point of the elbow plot. The perpendicular distance for each point was computed as follows:

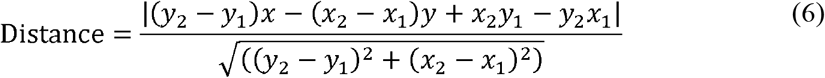

where (*x*_1_, *y*_1_) and (*x*_2_, *y*_2_) represent the first and last points on the straight line respectively, *x* denotes the max-normalized number of clusters (horizontal axis), and *y* denotes the max-normalized cost (vertical axis).

The P1acc-P2acc variable exhibited heavy right skew (skewness = 4.11). As a robustness check, we repeated the behavioral clustering after log-transforming acceleration difference prior to normalization. The cluster solutions were qualitatively identical.

For DTW, avatar velocities were computed for each 3-s epoch, normalized to [0, 1] within each epoch, and smoothed using a 20-point moving average. This normalization ensured that DTW captured similarity in temporal structure rather than absolute velocity. DTW was then computed with the MATLAB dtw function, which aligns the two signals and sums the Euclidean distances along the optimal warping path.

In the recording configuration, with the sampling rate and high-cutoff frequency equal, the BrainVision software automatically applied an anti-aliasing 450 Hz Butterworth low-pass filter (24 dB/octave), per the manufacturer.

#### S2.1 Adjusted circular correlation coefficient

Following the code provided by Zimmermann and colleagues [73], the adjusted CCorr is defined as:

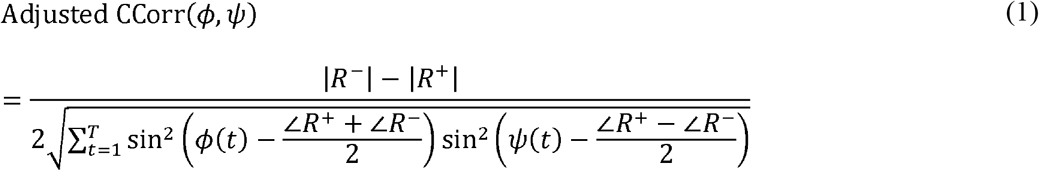

where R is the circular mean of phase signals:

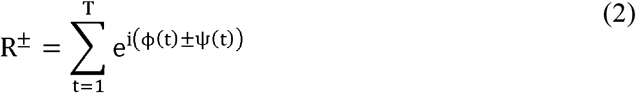

and *ϕ* and *ψ* are the phase signals of the neural signals *x* and *y* respectively. For example, *ϕ* is estimated with:

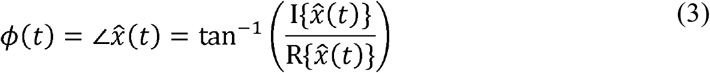

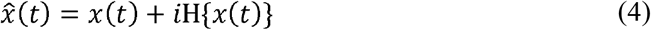

where H{*x*(*t*)} is the Hilbert transform of *x*(*t*), *x̂*(*t*) is the analytic signal of *x*(*t*), ℑ{*x̂*(*t*)} is the imaginary component, and ℜ{*x̂*(*t*)} is the real component of *x̂*(*t*). While the computed CCorr is bounded between [-1,1], we used the absolute value of CCorr, which bounds the coefficient between [0,1], as we were more interested in the magnitude of the synchronization rather than the direction of synchronization (whether in-phase or anti-phase).

#### S2.2 Dyad-shuffled null distribution and functional connectivity thresholding

Since participants were exposed to a similar external environment and shared similar stimuli in the PCE task space, we needed to discriminate between spurious connections due to task similarity and genuine connections resulting from the interaction process. Therefore, we created a null distribution for CCorr by shuffling the dyad numbers, such that 30*29 = 870 surrogate dyads were created. The trial numbers, epoch numbers, frequency bands and sensor numbers were not shuffled. This resulted in dyad-shuffled CCorr which constituted the null distribution, where each of the observed CCorr was tested against this null distribution. For example, for each dyad, trial, epoch, band, and sensor-sensor connection, the observed CCorr was ranked in the null distribution and thresholded at the 97.5th percentile rank in the null distribution. With 870 surrogates, there are 871 possible ranks, of which 22 exceed this threshold. Thus, under the null, a connection is expected to exceed the threshold by chance with probability 22/871 ≈ 0.025. Where epochs were rejected in a given surrogate pairing, fewer surrogate values were available for the corresponding epoch and connection, causing the empirical probability of exceeding the threshold by chance to deviate slightly from the nominal 0.025.

The dyad-shuffled surrogate procedure produced 30 × 18 × 20 × 7 binary FC matrices, retaining connections that exceeded the corresponding surrogate thresholds and were unlikely to be explained by shared task stimuli alone. To be more conservative, we further filtered out the weakest connections, retaining only the top 0.5% or top 0.1% of connections. This was done by first summing up all the FC matrices and thresholding at the 99.5th or the 99.9th percentile. This step resulted in one FC matrix for each frequency band, representing the connections that exceeded the surrogate threshold inter-brain connections during the PCE task.

In the same way as the behavioral variables, the un-thresholded FC matrices for the Alpha-Low and Alpha-High band were first flattened into vectors resulting in 30x18x20x4096 (DYADxTRIALxEPOCHxN) matrices. Similarly, the optimal number of clusters was determined by using the elbow method and the inter-brain FC matrices in the Alpha-Low and Alpha-High band were clustered using this optimal number of clusters with the number of replicates set to 20. After labeling each epoch with a cluster number, we then pooled and summed the FC matrices for each cluster number. Finally, these clusters were thresholded separately at the 99.5th percentile, retaining only the top 0.5% of the strongest connections.

#### S3.1 Sensitivity analyses of Spearman correlations across levels of aggregation

To address the non-independence of epochs nested within trials within dyads, each Spearman correlation was repeated after aggregation to the trial level (*n* = 538, the trials retaining at least one epoch after artefact rejection) and to the dyad level (*n* = 30; S7 Fig), with the complete numerical results provided in tabs1_results.csv in the code repository. The ROI connections were those retained in Fig 3C, eight for Alpha-Low and six for Alpha-High, and the same Benjamini-Hochberg correction was applied within each band and level.

The correlation coefficients increased as the unit of analysis became coarser, from approximately .03 to .05 at the epoch level to .20 to .39 at the dyad level, while statistical significance decreased as *n* fell from 10,440 to 30. Coefficients estimated from fewer observations can be larger in absolute value even under the null hypothesis, because the sampling variability of a correlation increases as the number of units decreases. At the dyad level (n = 30), the largest coefficient observed across this family of tests was approximately .39, comparable in magnitude to the largest coefficient expected under the null. Thus, the increase in coefficient magnitude with aggregation does not provide evidence for a stronger association at the dyad level, and the aggregated analyses neither support nor rule out such an association. Of the 27 cells surviving correction, all 27 were significant at the epoch level, and none at the trial or dyad level. At the dyad level the same two variables were the largest in both bands. Avatar-shadow distance ranged from *ρ* -.390 to -.335 in Alpha-Low and from *ρ* -.200 to -.109 in Alpha-High. Inter-avatar distance ranged from *ρ* +.227 to +.330 in Alpha-Low and from *ρ* +.204 to +.272 in Alpha-High.

#### S3.2. Within-trial permutation test for the Spearman correlations

The aggregation analysis in S3.1 reduces the number of observations and therefore statistical power. To retain all epochs while accounting for their dependence, we constructed an empirical null distribution for each Spearman correlation using the same procedure as for the chi-square statistic in S4.2. Within each trial, the seven behavioral variables were jointly and circularly shifted by a randomly selected offset, with values wrapped around, while the inter-brain connection values were held fixed. Each shift preserves the distribution of each behavioral variable and its temporal run structure within each trial, as well as the trial and dyad membership of every observation, while breaking the epoch-by-epoch alignment between behavior and inter-brain coupling. The correlation was recomputed after each of 10,000 shifts, and a two-sided permutation *p*-value was calculated as the proportion of shifts for which the absolute correlation equalled or exceeded the observed value. Benjamini-Hochberg correction was then applied to the permutation *p*-values within each band.

Under this null, none of the 14 Alpha-Low cells or 13 Alpha-High cells that were significant under the parametric test remained significant after correction. The largest Alpha-Low correlation, between inter-avatar distance and the RF-LP connection (*ρ* = .049), had a parametric adjusted *p*-value below .001 but a permutation adjusted *p*-value of .40. The largest Alpha-High correlation, between inter-avatar distance and the RT-LT connection (*ρ* = .038), had a parametric adjusted *p*-value of .004 but a permutation adjusted *p*-value of .31. The smallest permutation adjusted *p*-value in either band was .28. Thus, none of the epoch-level associations was robust to the dependence-aware null, consistent with the loss of significance under aggregation in S3.1 and with the cluster-level results in S4.

#### S4. Dyad-level analysis of behavioral to inter-brain associations

Adjusted Pearson residuals can be computed by:

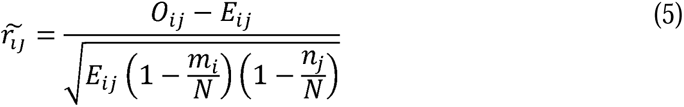

where *O_ij_* represents the observed count in cell (*i*, *j*), *E_ij_* was the expected count under the null hypothesis of independence, *m_i_* and *n_j_* are the marginal sums for row *i* and column *j*, respectively, and *N* is the grand total count of all observed frequencies. Adjusted Pearson residual takes into consideration the row and column totals, thus providing a more precise measure of deviations from the expected counts. Furthermore, the asymptotic normality for Pearson residuals is not always observed in real data and using Pearson residuals alone can underestimate sample variability [10]. In contrast, adjusted Pearson residuals are asymptotically normal-distributed, providing more accurate statistical inferences than using Pearson residuals. These residuals can be interpreted as effect sizes. Larger values can be interpreted as stronger effects, positive values represent a positive association, and negative values represent a negative association between a category in the rows and a category in the columns. In particular, an association means that there is a relationship between the two categories, for example inter-brain cluster 1 and behavioral cluster 3, such that one category is more (positive) or less (negative) likely to occur with the other.

Cramér’s V was computed as the effect size, with a 95% confidence interval obtained by resampling epochs with replacement from the observed table 10,000 times using the rmultinom function in R and taking the 2.5th and 97.5th percentiles of the resulting distribution of V.

The chi-square test of independence reported in the main text treats all 10,440 epochs as independent observations. However, epochs are nested within trials, and trials within dyads, such that a dyad that spends much of its time in one behavioral mode and one inter-brain mode contributes repeated observations to the same apparent association. Because the chi-square statistic is sensitive to sample size, treating these non-independent observations as independent can yield anti-conservative significance estimates, potentially allowing a weak association to reach the conventional significance threshold. This interpretation is consistent with the small Cramér’s V values reported in the main text. We addressed this issue in two ways: one that explicitly models the dependency structure and one that avoids the corresponding reference distribution.

The two approaches yielded consistent conclusions. The permutation test provided a more conservative assessment, as the circular shifts preserve the temporal structure within trials, which is not explicitly modeled by the random-intercept structure of the mixed-effects model. We therefore conclude that the association between behavioral and inter-brain clustering is not supported when the nesting of epochs within trials and dyads is taken into account.

##### S4.1. Mixed-effects logistic regression

We re-examined the same association by fitting a mixed-effects logistic regression to the epoch-level data:

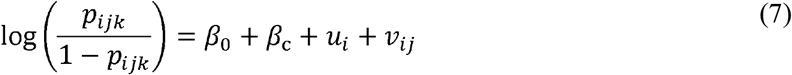

where *p_ijk_* is the probability that epoch *k* of trial *j* in dyad *i* was assigned to an inter-brain cluster other than the dominant one, *c* is the behavioral cluster of that epoch entered as a four-level categorical predictor with cluster 1 as reference, and *u_i_* and *v_ij_* are random intercepts for dyad and for trial within dyad, with *u_i_* ∼ N(0, σ^2^D) and *v_ij_* ∼ N(0, σ^2^T). These random effects allow the baseline probability to vary across dyads and trials, such that clustering at these levels is accounted for rather than contributing to the fixed effect. The model was fitted in MATLAB R2022b using *fitglme*, as minority ∼ 1 + beh + (1|dyad) + (1|dyadtrial), using maximum likelihood with a binomial distribution, logit link, and Laplace approximation. The behavioral effect was tested using a likelihood-ratio test comparing this model with an otherwise identical model excluding the behavioral-cluster term, with three degrees of freedom. The intraclass correlation coefficient was computed as (*σ*^2^D + *σ*^2^T) / (*σ*^2^D + *σ*^2^_T + π^2^/3).

The behavioral effect was not significant in either band (Alpha-Low, □^2^ (3) = 7.65, *p* = .054; Alpha-High, □^2^ (3) = 6.90, *p* = .075). The random effects were substantial. On the log-odds scale, the *SD* of the dyad-level intercept was 1.98 in Alpha-Low and 2.26 in Alpha-High, while the *SD* for trials nested within dyads was 0.47 and 0.52, respectively. The corresponding latent-scale intraclass correlations were .56 and .62, indicating substantial clustering of epochs by dyad and trial, with more than half of the latent-scale variance attributable to these grouping factors before behavioral cluster was considered.

##### S4.2. Permutation null for the chi-square

The mixed-effects model requires collapsing the three inter-brain clusters into a binary contrast. To retain the full contingency table, we also constructed an empirical null distribution for the chi-square statistic itself. Within each trial, the behavioral label sequence was circularly shifted by a randomly selected offset and wrapped around, while the inter-brain labels were held fixed, and the chi-square statistic was recomputed. This procedure was repeated 10,000 times. Each shift preserves the behavioral and inter-brain cluster composition of every dyad and trial and largely preserves the temporal run structure of both label streams, as a circular shift disrupts a run at only one boundary. The procedure therefore disrupts the temporal alignment between the two label streams within each trial while preserving their marginal distributions and within-trial structure.

Under the empirical null, the association was not significant (Alpha-Low, *p* = .307; Alpha-High, *p* = .433), in contrast to the nominal *p*-values obtained from the theoretical chi-square distribution (*p* = .019 and .0498, respectively). The difference between the two reference distributions is evident from the null distribution itself: for Alpha-Low, its mean was 12.88 and its 95th percentile was 23.69, compared with 6.00 and 12.59, respectively, for the χ^2^(6) distribution assumed by the standard test. The observed statistic of 15.17 therefore lies only slightly above the mean of the empirical null distribution.

The same circular shifts were used to construct empirical null distributions for each adjusted Pearson residual. No cell survived Benjamini-Hochberg correction under either reference distribution, although the permutation *p*-values were consistently larger. The smallest permutation *p*-values were .255 compared with .060 in Alpha-Low and .436 compared with .113 in Alpha-High. Tables 3 and 4 report the Benjamini-Hochberg-corrected *p*-values from both the theoretical chi-square reference distribution (*p*) and the within-trial permutation null (*p*_perm_).

### Supplementary Figures

**S1 Fig.**
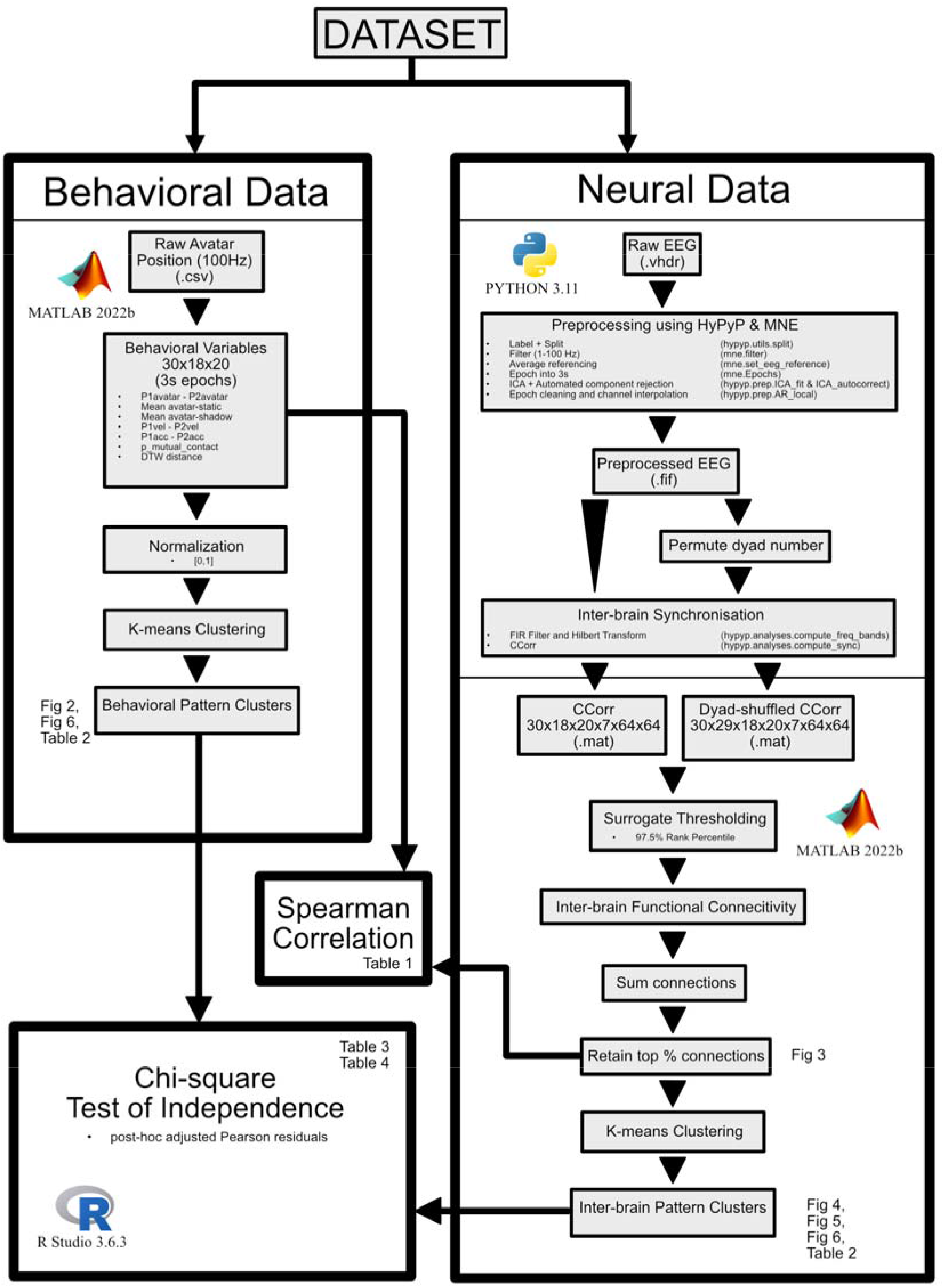
Overview of the analysis plan. Behavioral and inter-brain data were first analysed separately. Spearman correlation was computed between regions of interest in the Alpha-Low (7.5-10 Hz) and Alpha-High (10-12.5 Hz) bands and the behavioral variables. A contingency table of the behavioral and inter-brain clusters was created to investigate the association between behavioral and inter-brain clusters using chi-square test. The figure also pointed to where the figures and tables were created from. CCorr = circular correlation; DTW = dynamic time warping; EEG = electroencephalography.

**S2 Fig.**
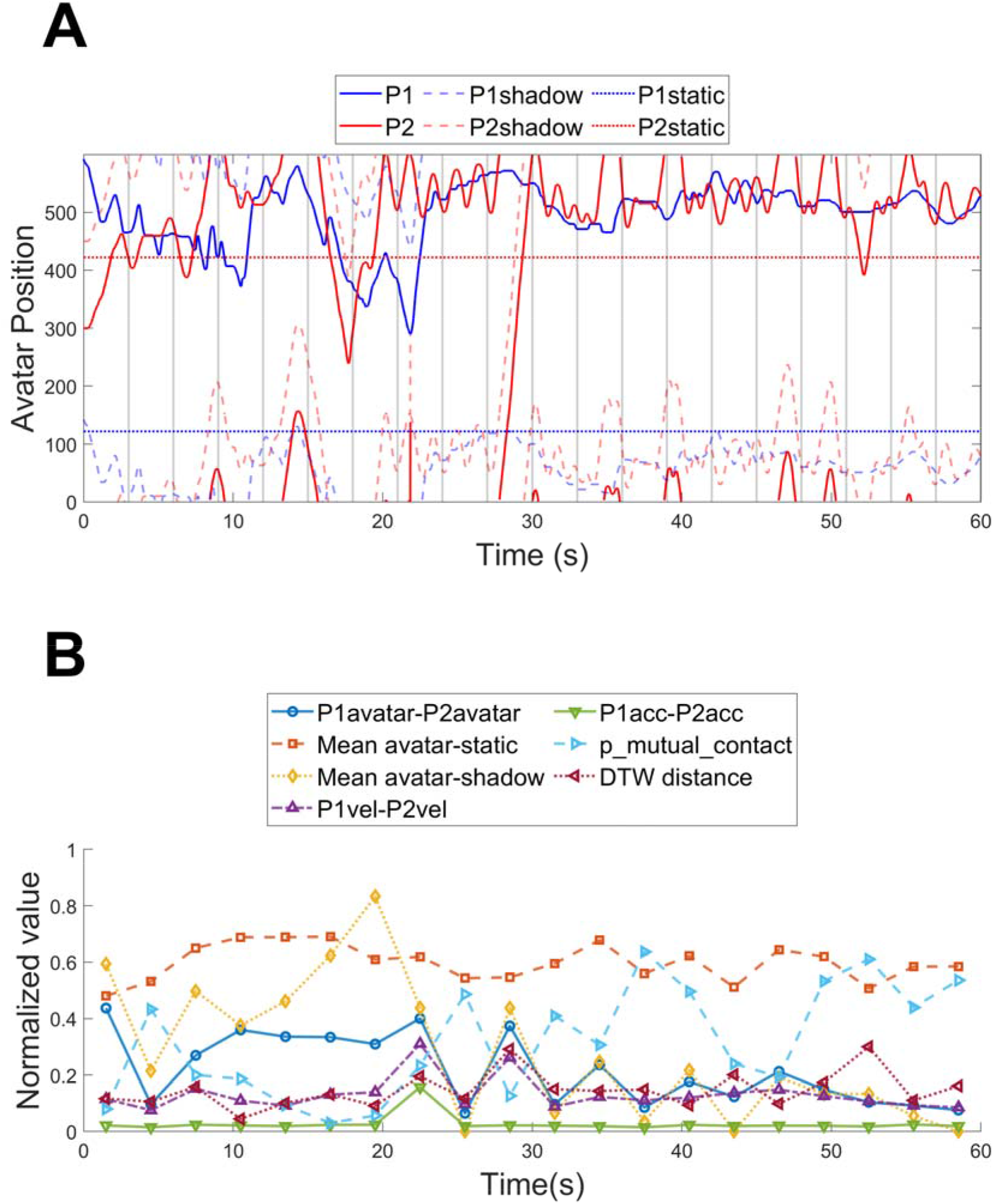
Example plot of behavioral data. (A) Avatar, shadow and static object positions of participant 1 and participant 2. Grey vertical lines mark 3-second windows. P1= participant 1; P2 = participant 2; avatar = avatar position; shadow = shadow position; static = static object position. (B) Derived behavioral variables. DTW = dynamic time warping.

**S3 Fig.**
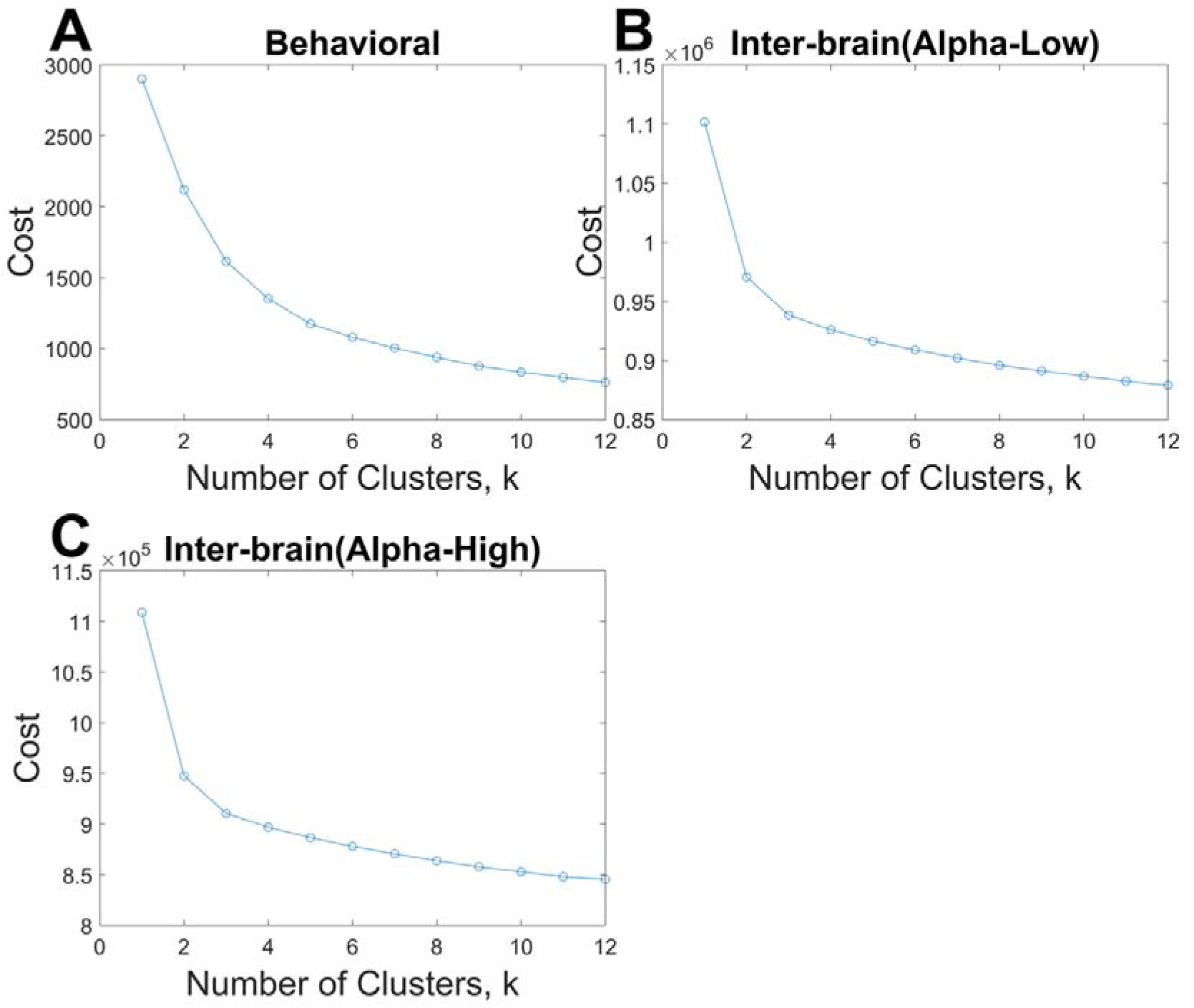
Elbow plots for the detection of the optimal number of clusters, **k.** (A) Cost and k for the clustering of behavioral variables. The calculation of the optimal k could be understood as drawing a straight line from the first to the last point of the curve. The perpendicular distance from this line to the elbow curve was then calculated for each k. The optimal k was the k at which the perpendicular distance was maximum. Cost and k for the clustering of inter-brain patterns in the (B) Alpha-Low and (C) Alpha-High bands.

**S4 Fig.**
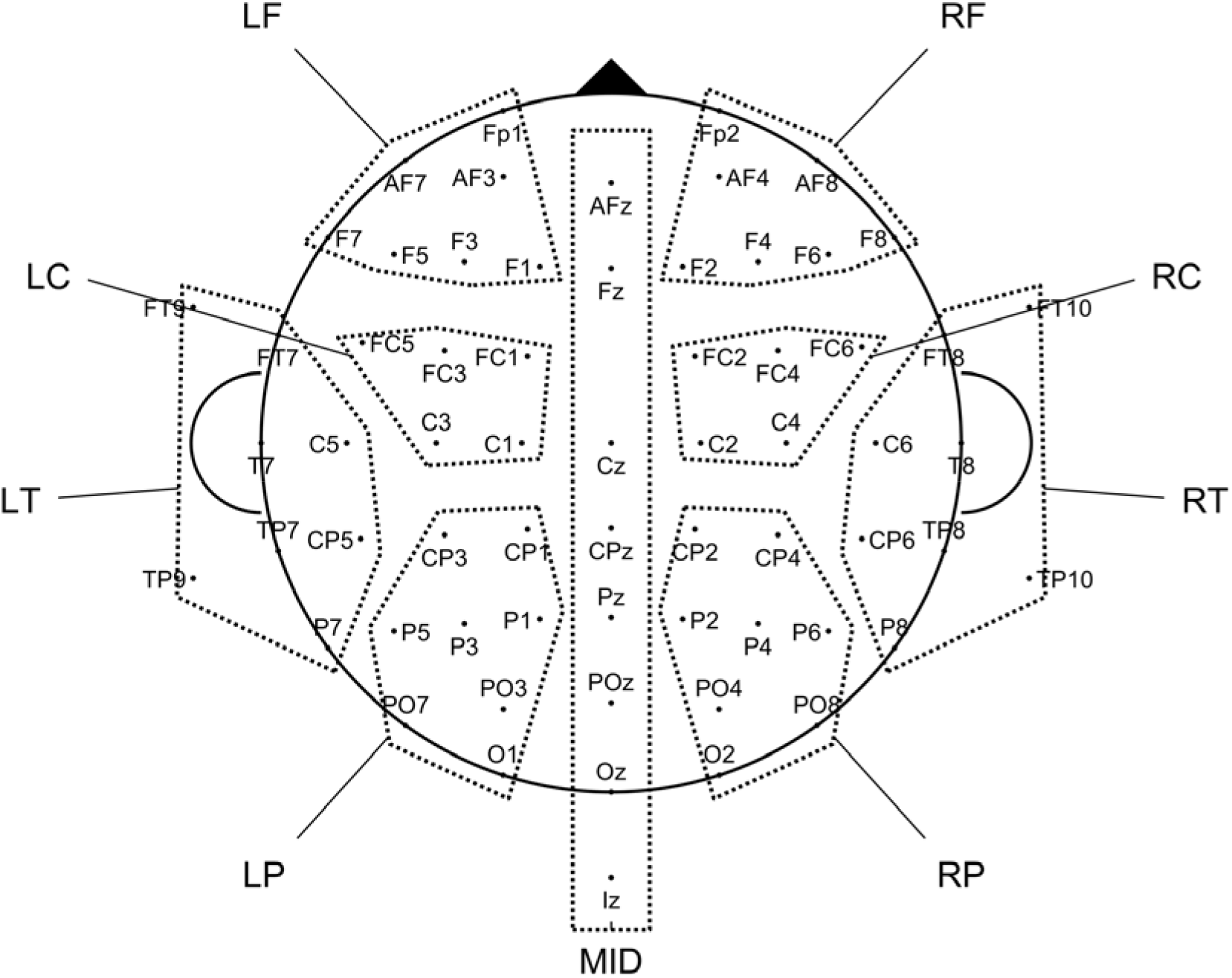
Regions of interest. Head plot showing the grouping of all 64 sensors into 9 regions of interest (ROIs). RF = right frontal; LF = left frontal; RC = right central; LC = left central; RT = right temporal; LT = left temporal; RP = right parietal; LP = left parietal; MID = midline.

**S5 Fig.**
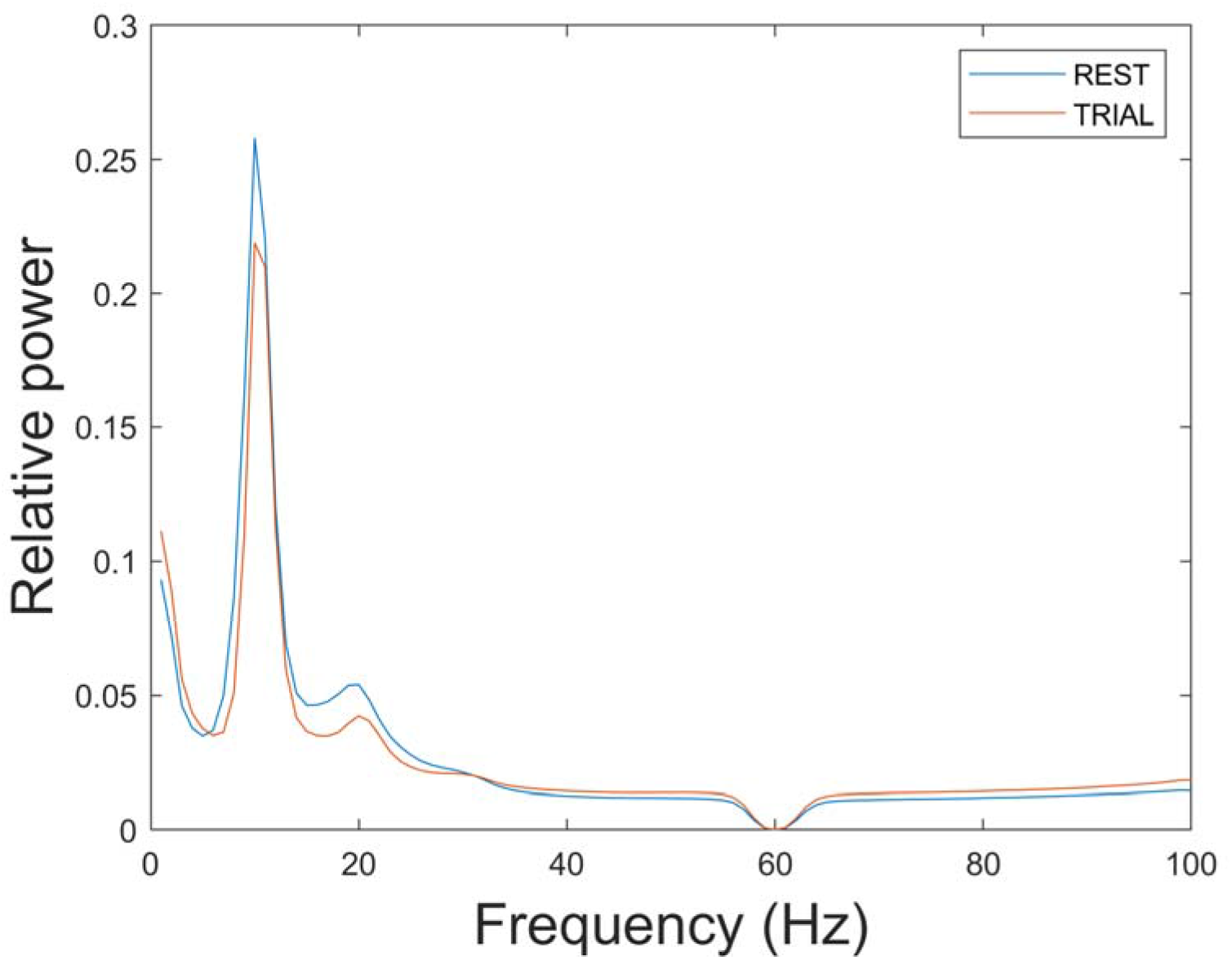
Relative Welch power spectral density for the 60 participants in the Perceptual Crossing Experiment, averaged over all epochs for resting phase and experimental trials. REST= resting phase; TRIAL = experimental trials.

**S6 Fig.**
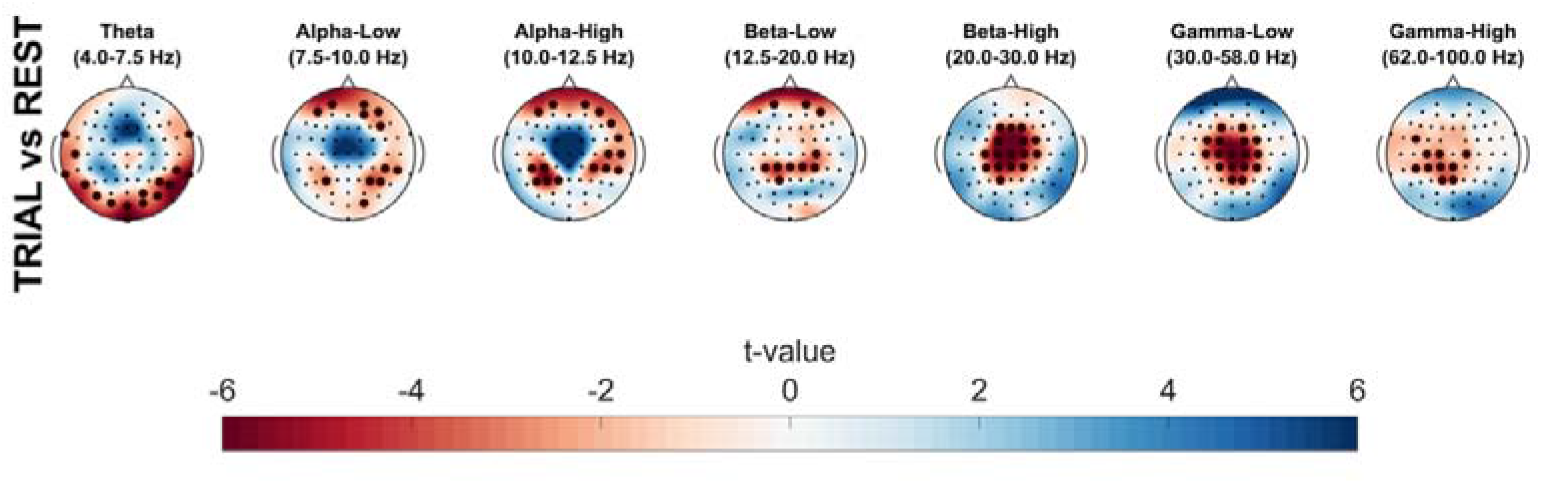
T-statistics of the differences in relativeWelch power spectral density between resting phase and experimental trials. Asterisks (*) marked significant differences at a sensor and frequency band, tested using the MNE permutation cluster test. Dark red, TRIAL < REST; Dark blue, TRIAL > REST. REST= resting phase; TRIAL = experimental trials.

**S7 Fig.**
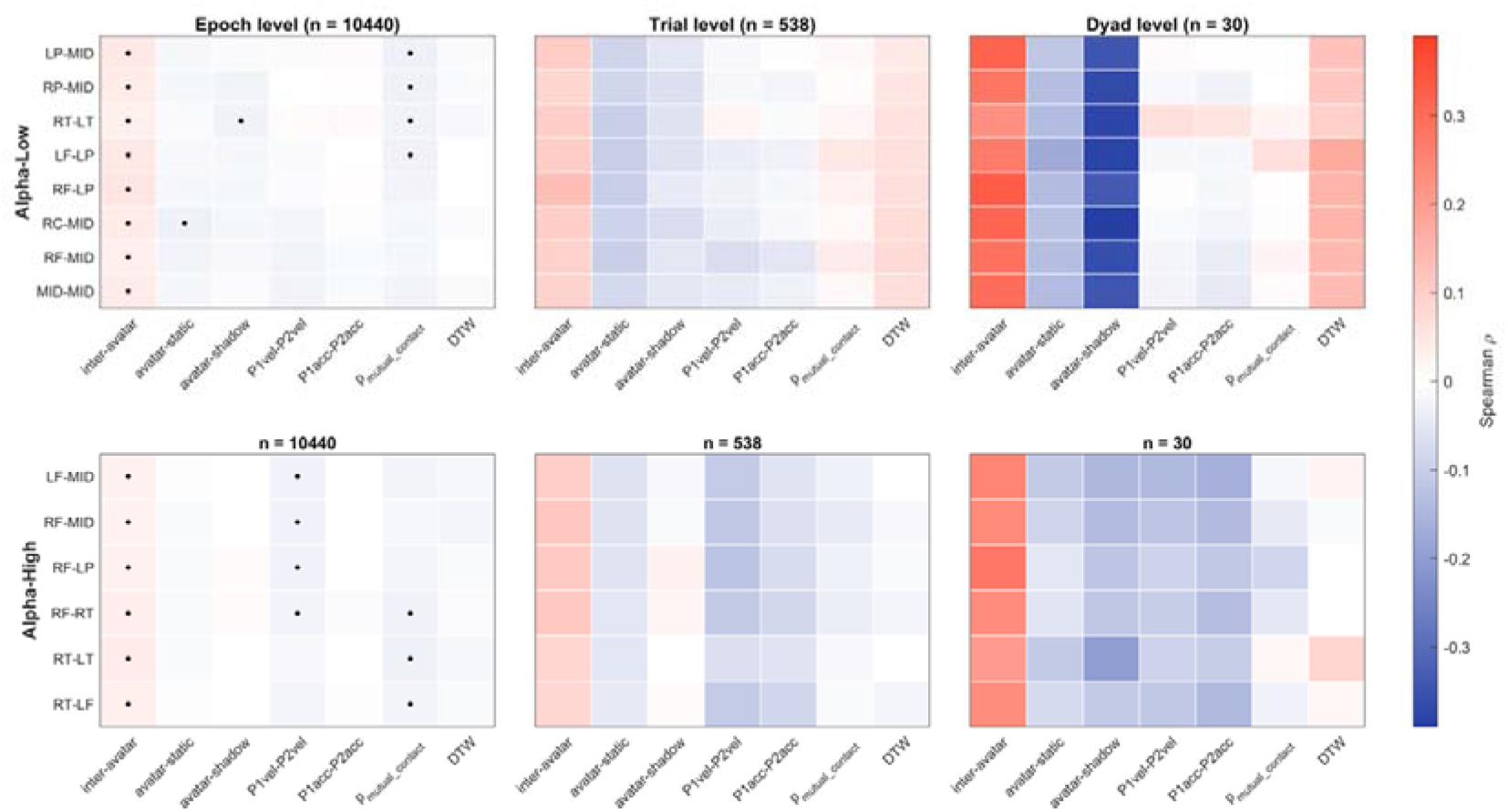
Spearman correlations between behavioral variables and inter-brain connections at three levels of aggregation. Rows are the ROI-ROI connections retained in Fig 3C, eight for Alpha-Low and six for Alpha-High, and columns are the seven behavioral variables. Panels show the epoch level (*n* = 10,440), the trial level (*n* = 538) and the dyad level (*n* = 30). Cell color gives the Spearman correlation coefficient, and a dot marks each cells surviving Benjamini-Hochberg correction within each band and level. This figure shows all connection by variable combinations, including those not listed in Table 1. Complete numerical results for all 294 tests are provided in tabs1_results.csv in the code repository.

